# Hazard characterization of *Alternaria* toxins – filling data gaps on *in vitro* genotoxicity

**DOI:** 10.64898/2026.07.08.737172

**Authors:** Anne-Cathrin Behr, Ariane Vettorazzi, Camille Streel, Birgit Mertens, Roel Antonissen, Beatriz Guerreiro, Célia Ventura, Rita Sofia Vilela, Matjaz Novak, Bojana Zegura, Fabian Reith, Lennart Oltmanns, Victor Prisyazhnoy, Roderich D. Süssmuth, Maria João Silva, Henriqueta Louro, Doris Marko

## Abstract

*Alternaria* toxins are naturally occurring food contaminants with limited and often inconsistent genotoxicity and mutagenicity data. Within the European Partnership for the Assessment of Risks from Chemicals (PARC), an OECD-aligned *in vitro* testing strategy was applied to fill existing data gaps and to characterize the genotoxic potential of major *Alternaria* toxins using high-purity test materials. Mutagenicity was assessed using bacterial reverse mutation test (OECD TG 471) and SOS/umu assay, while chromosomal damage was assessed using the *in vitro* micronucleus (MN) assay (OECD TG 487) in TK6 and HepG2 cells, complemented by fluorescence *in situ* hybridization (FISH) and γH2AX assay in HepaRG cells.

Alternariol (AOH), alternariol monomethyl ether (AME), and altertoxin-I (ATX-I) showed clear mutagenicity in bacteria, whereas altenuene (ALT), tenuazonic acid (TeA), and tentoxin (TEN) were negative under the tested conditions. In mammalian cells, AOH, AME, and ATX-I induced MN formation in TK6 cells at concentrations ≥5.5 µM, ≥2.5 µM, and ≥0.21 µM, respectively, with FISH analysis supporting a clastogenic mode of action. In HepG2 cells, all tested toxins induced chromosomal damage, with effect threshold ranging from ≥6.25 µM (AOH) to ≥50 µM (TeA). γH2AX induction confirmed DNA damage for AOH and ATX-I, and at higher concentrations for TeA (1000 µM).

Overall, the data indicate clear *in vitro* genotoxic potential for AOH, AME, and ATX-I and provide evidence of chromosomal damage for ALT, TEN, and TeA, thereby reducing critical data gaps for hazard assessment.

## 1. Introduction

The European Partnership for the Assessment of Risks from Chemicals (PARC) is focused on overcoming the major challenges in hazard assessment to protect human health and the environment. Following a prioritization workshop and consultations with partners from national and European agencies involved in hazard and risk assessment, one substance group that has been selected for as a priority for hazard assessment was natural toxins, with a focus on emerging mycotoxins such as enniatins (produced by *Fusarium* fungi) and *Alternaria* toxins.

Fungi of the genus *Alternaria* are saprophytes and represent ubiquitous plant pathogens. *Alternaria* spp. can infest a broad spectrum of substrates and is known to grow under a wide range of environmental conditions. So far, more than 250 secondary metabolites out of different structural classes formed by *Alternaria* spp. have been identified (Zhao et al. 2023), but only a few of these secondary metabolites are commercially available in sufficient purity to allow hazard characterization. Important structural classes comprise dibenzo-α-pyrones, perylene quinones, tenuazonic acids and other miscellaneous structures such as tentoxin (Fig 1).

**Fig 1:**
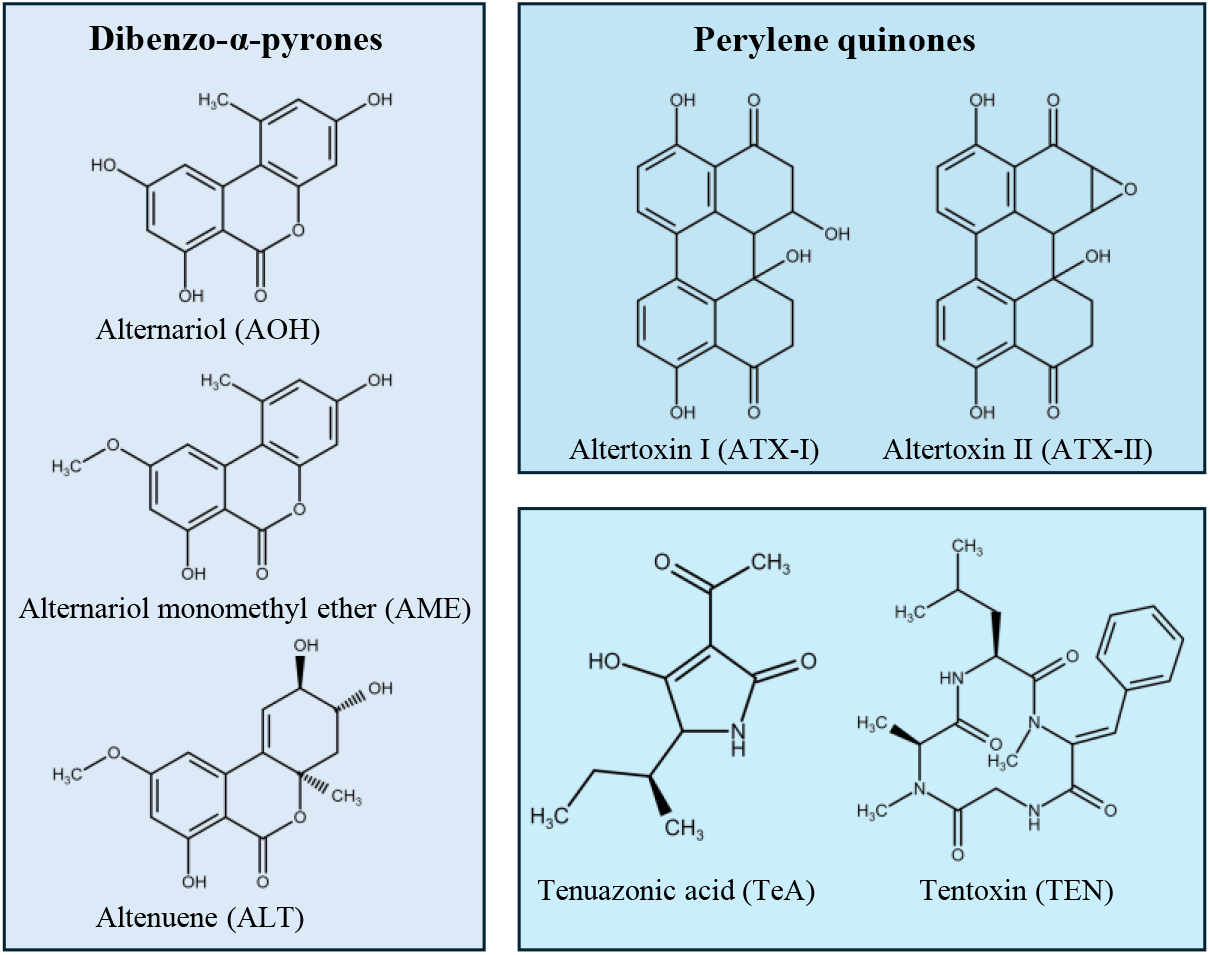
Structures of *Alternaria* toxins

Depending on the infesting strain, available substrate and growth conditions, the formed spectrum of secondary metabolites vary substantially. In food, most data are available on the occurrence of the dibenzo-α-pyrones alternariol (AOH) and its monomethyl ether (AME), tenuazonic acid (TeA) and tentoxin (TEN) (EFSA 2011b; EFSA 2016). Several *Alternaria* toxins have been reported to exhibit genotoxic and mutagenic potential but through different modes of action. The genotoxic properties of TeA and TEN remains to be clarified. AOH and AME have been characterized as inhibitors of topoisomerase II (El-Sayed et al. 2026; Fehr et al. 2009), thus interfering with the regulation of DNA topology. Little is known so far about the genotoxic potential of the structurally related altenuene (ALT). The class of perylene quinones still represents a challenge concerning occurrence and hazard characterization. Previous studies indicate potent genotoxic properties especially of epoxide-bearing analogues such as altertoxin II (ATX-II; Fig 1) which are likely to form DNA adducts (Soukup et al. 2020). However, the epoxide-bearing analogues are not commercially available. Furthermore, due to the reactivity of the epoxide moiety, the question arises whether these compounds occur in food. ATX-II has been identified in naturally infected as well as inoculated apples (Pavicich et al. 2025; Puntscher et al. 2020) but the fate of the compound during food processing and digestion remains to be elucidated. In a gastromimetic study, the level of free ATX-II decreased substantially until the intestinal phase (Call et al. 2026). Application of ATX-II to enzymatically active tomato material resulted in a quick loss of free ATX-II and the formation of the less reactive altertoxin-I (ATX-I; Fig 1) (Puntscher et al. 2019b). In contrast to the epoxide-bearing analogues, ATX-I is commercially available in sufficient purity for hazard characterization and was included in the present study.

So far, *Alternaria* toxins are still considered as emerging mycotoxins. Inconsistencies across reported *in vitro* studies highlight the need for additional testing, preferably following OECD guidelines, to address the current data gaps and support risk assessment. Overall, further experiments with higher-purity compounds and adequately selected test strategies are needed. The aim of the present study was to engage multiple PARC partners to fill the data gaps using established toxicological tests, in line with OECD guidelines, complemented by supporting sensitive methods to ensure accurate assessment of the genotoxic properties of *Alternaria* toxins available in sufficient purity. For allowing an integrated assessment, all endpoints selected for the studies were developed using the same batch of high-quality toxins and standardized procedures in preparing the toxins for testing under each assay. This manuscript describes the preparation of the toxins and the *in vitro* genotoxicity testing that was performed. Overall interpretation of the data is provided considering the OECD criteria available for positive, negative or equivocal results.

## 2. Material and Methods

### 2.1. Mycotoxins

Alternariol (AOH) was synthesized according to Won et al. (Won et al. 2015). The convergent synthesis featured a formylated aryl bromide and a desymmetrized aryl borane. Both were connected during a Pd-mediated Suzuki coupling. The final ring closure of the oxidized biaryl was performed with an ester formation during demethylation of all remaining alcohols. The product was isolated using reverse phase column chromatography and the identity of the material was confirmed by analytical HPLC-ESI-MS(MS) and 1H-NMR spectroscopy. The convergent synthesis of alternariol-9-methyl ether (AME) was carried out according to Mikula et al. (Mikula et al. 2013). The building blocks were prepared from orcinol and 3,5-dimethoxyaniline respectively and coupled using diethyl chlorophosphate (DECP). The benzocoumarin core structure was established by intramolecular cyclization of the iodoresorcylic acid phenyl ester. Selective demethylation of alternariol 2,4-dimethyl ether with AlCl3/NaI yielded AME which was isolated using reverse phase column chromatography. The identity of the material was confirmed by analytical HPLC-ESI-MS(MS) and 1H NMR spectroscopy. The identity of the synthesized compounds was checked by liquid chromatography (Vanquish UHPLC, Thermo Fisher Scientific) coupled to a high-resolution mass spectrometer (timsTOF flex, Bruker) with a mass accuracy of < 2 ppm. Furthermore, the purity was estimated by the absence of significant peaks in the base peak chromatogram in the respective retention time window (both in negative and positive ionization mode, target peak > 98% of the peak area) and the UV signal at 220 nm (target peak > 97% of the peak area). The presence of possible isomers as byproducts of the synthesis procedure could be ruled out this way.

Tenuazonic acid (TeA) was purchased from SantaCruz (Heidelberg, Germany), and altertoxin-I (ATX-I), (−)-altenuene (ALT) and tentoxin (TEN) from Biomol (Hamburg, Germany). To allow for data comparability while reducing uncertainties related to the test substances, the assays were carried out using te same batch of each mycotoxin.

Upon arrival, dry compounds were stored at −80°C. The toxins were dissolved in dimethyl sulfoxide (DMSO; Merck, Darmstadt, Germany) at the maximum concentration indicated in Table 1. For the SOS/umu assay, an additional, higher concentration (2x the maximum concentration) was also tested for ALT, TeA and TEN. For the *in vitro* MN assay in TK6 cells, stock solutions with higher concentration were also made for AME (25 mM), ALT (20 mM) and TEN (20 mM) to test an additional higher concentration. The stock solutions were vortexed for 1 min and subsequently sonicated for 10 min in a water bath. Small aliquots were prepared in amber glass vials to protect the compounds from light and to avoid more than three freeze-thaw cycles. The stock solutions were stored at −20°C. The final concentrations employed in each assay were determined by the solubility of the product under the assay-specific conditions and are reported in the corresponding section for each assay.

**Table 1:**
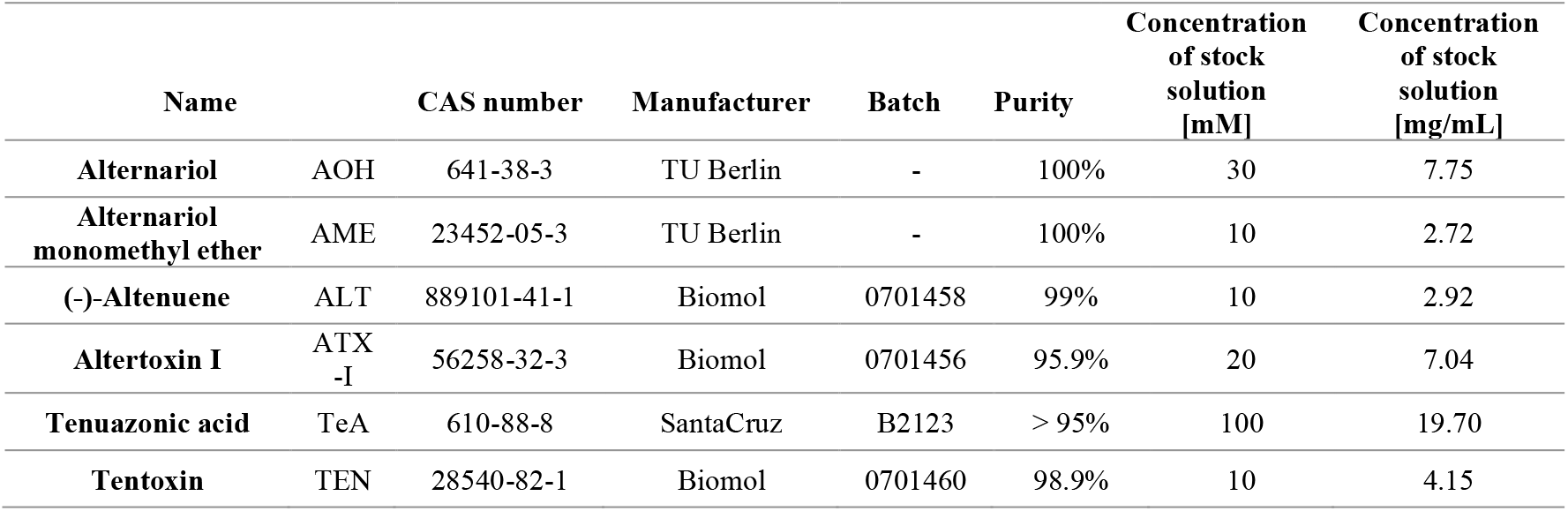
Information on test items and the concentration of stock solutions.

### 2.2. Bacterial Reverse Mutation test

#### Bacteria

*Salmonella typhimurium* (*S.Typhimurium*) strains TA97a (71-097aL, 4710D), TA98 (71-098L, 4599D), TA100 (71-100L, 4598D), TA102 (71-102L, 4604D), and TA1535 (71-1535L, 4602D) were obtained from Trinova Biochem GmbH, Germany. The test batches of bacterial culture are stored in nutrient broth with 10% DMSO until use (Merck, Germany) as a cryopreservative at −80 ± 15°C. The strains are routinely checked to confirm their histidine-requirement, the presence of *rfa* (crystal violet sensitivity), *uvrB*-sensitivity and the R-factor plasmid pKM101 (TA97a, TA98, TA100, TA102; ampicillin resistance) and presence of R-factor plasmid pAQ1 (TA102; tetracycline resistance) and the number of spontaneous revertants. TA1535 does not contain the R-factor plasmid (ampicillin non-resistant strain). Each tester bacterial strain reverts spontaneously at a frequency that is characteristic for each strain.

#### Ames assay

The Ames assay was performed according to OECD Test Guideline (TG) 471 under GLP-like conditions, with minor modification (OECD 2020). The highest tested concentrations were 0.1, 0.272, 0.292, 0.01, 2.5 and 0.415 mg/plate for AOH, AME, ALT, ATX-I, TeA, and TEN corresponding to 1, 2.72, 2.92, 0.1, 25 and 4.15 mg/mL in DMSO, respectively. These concentrations were selected based on available solubility data (Table 1) or literature reports on toxicity and mutagenic responses. The assay was performed using the plate incorporation method, both without and with metabolic activation system (post-mitochondrial rat liver fraction S9, Trinova Biochem GmbH, Germany).

In addition to mutagenicity testing, also solubility in the solvent, phosphate buffer, overlay agar, and on the plates were tested. Into separate tubes of 2 mL overlay agar supplemented with histidine-biotine (final concentration 0.05 mM) solution, 0.1 mL of the toxin at different concentrations and 0.5 mL phosphate buffer or 10% S9 mixture (cofactor-supplemented post-mitochondrial fraction) were added. The prepared mixtures were inoculated with 0.1 mL of the corresponding bacterial culture for each of five tester strains, except for TeA which was tested only in strains TA98 and TA100. Each mixture was poured on the surface of the minimal glucose agar plates (Minimal E plates) in triplicates and incubated for two (*S.Typhimurium* TA97a, TA100, TA102, TA1535) or three days (*S.Typhimurium* TA98). In parallel, testing with negative control (sterile milliQ-Water), solvent control (DMSO) and tester strain specific reference materials (Sigma Aldrich, Germany) was conducted.

#### Data analysis

After the incubation period, the revertant colonies on each plate were counted manually and data are presented as the number of revertants / plate per concentration of the test item, negative and reference controls, along with mean values and the standard deviations (SD). The induction factor (IF) was calculated by dividing the mean number of revertants obtained when testing the toxin or reference materials with the mean number of revertant colonies obtained with the solvent control. Additionally, the background lawn quality (as an indicator of cytotoxicity) was assessed for signs of toxicity, and potential precipitation of the test item was also checked.

The toxin is considered mutagenic if at least 2-fold increase (IF ≥ 2, for strains TA97a, TA98, TA100 and TA102) or 3-fold increase (IF ≥ 3 for strain TA1535) in the mean revertants per plate over the mean solvent revertants in at least one bacterial strain without or with metabolic activation is observed at least at the highest test item concentration, where no sign of cytotoxicity is observed (statistical significance is not the determining factor for positive result).

Criteria for bacterial background lawn quality relative to the solvent control plate evaluated by using inverted light microscope:

1) *Normal*; distinguished by a healthy bacterial background lawn.
2) *Slightly reduced*; distinguished by a noticeable thinning of the bacterial background lawn and possibly a slight increase / decrease in the size of the microcolonies compared to the solvent control plate.
3) *Moderately reduced*; distinguished by a marked thinning of the bacterial lawn that may result in a pronounced increase in the size of microcolonies compared to the solvent control plate.
4) *Extremely reduced*; distinguished by an extreme thinning of the bacterial background lawn that may result in an increase in the size of the microcolonies compared to the solvent control plate such that the microcolony lawn is visible to the unaided eye as isolated colonies – pinpoint colonies.
5) *Absent*; distinguished by a complete lack of any bacterial background lawn over greater than or equal to 90 % of the plate.
6) *Obscured by precipitate*; the background bacterial lawn cannot be accurately evaluated due to microscopic test substance precipitate.

The concentration of the test item was considered to be cytotoxic when the microcolony lawn is classified as; *moderately reduced, extremely reduced, absent* or a > 50 % reduction (IF < 0.5) in the mean number of revertants per plate as compared to the mean vehicle value obtained.

### 2.3. SOS/umu Assay

#### Bacteria

The genetically modified *S.Typhimurium* 1535/pSK1002 used in the SOS/umu test was purchased from the German Collection of microorganisms and Cell cultures (DSMZ; 9274, Braunschweig, Germany). The bacterium contains a plasmid in which two genes are fused, one involved in DNA repair and the other accountable for β-galactosidase activity. Therefore, SOS repair activity was monitored through β-galactosidase induction activity determined with spectrophotometric measures.

#### SOS/umu Assay

The SOS/umu test was performed according to the method used by Alonso-Jauregui et al. (2021) with some modifications. The bacteria were incubated overnight at 37°C in 100 mL TGA medium (Bacto tryptone, Gibco, USA; Glucose, PanReaC, Spain; Sodium chloride, Sigma-Aldrich, Germany) supplemented with ampicillin (50 µg/mL), with slight orbital shaking (155 rpm) from 15 to 17h until an optimal orbital density (OD₆₀₀ from 0.5 to 1.5) was reached. Then, the overnight culture was diluted with fresh TGA medium (not supplemented with ampicillin) and incubated for 2h at 37°C with orbital shaking (155 rpm) in order to obtain a log-phase bacterial growth culture (OD₆₀₀ from 0.15 to 0.4). The test was performed in the absence (phosphate-buffered saline; PBS) and presence of an external metabolic activation system (S9 from Sprague Dawley rat liver S9, PB/BNF induced) from Moltox (Boone, USA). The S9 mix (10%) was prepared with phosphate buffer (NaH_2_PO_4_xH_2_O, Na_2_HPO_4_x2H_2_O), glucose-6-phosphate and NADP solutions, of which compounds were obtained from Sigma-Aldrich (Darmstadt, Germany) and saline solution ingredients (MgCl_2_x6H_2_O, KCl) from PanReac AppliChem (Barcelona, Spain). Afterwards, the S9 mix was filtered through a 0.45 µm filter. In each test, negative and positive controls were included. The positive control stock solutions were prepared in DMSO: at 500 µg/mL (corresponding to 12.5 µg/mL in 96-well plate) for 2-aminoanthracene (2-AA) (Sigma-Aldrich, Germany) and at 100 µg/mL (2.5 µg/mL) for 4-nitroquinoline-n-oxide (4-NQO) (Sigma-Aldrich, China). 4-NQO was the positive control without metabolic activation and 2-AA with metabolic activation (S9).

The test procedure was as follows: first, mycotoxins were dissolved in DMSO at the double of their respective maximum concentrations (Table 1) for ALT, TeA and TEN or at the maximum soluble concentrations in the assay conditions (AOH, AME, ATX-I). The final concentration in the 96-well plate corresponds to a 40-fold dilution from the DMSO stock solutions.

For the experiment, 11 serial half dilutions in DMSO of mycotoxins and positive controls were prepared in a 96-well plate (plate A). The final volume in each well was 10 µL. The wells on the last row of the plate contained the negative control (DMSO). Afterwards, 70 µL water was added to each well. At this point, each well was checked in order to detect any precipitation of the mycotoxins. Then, the S9 mix was prepared as mentioned before. Thereafter, in another 96-well plate (plate B), 10 µL S9 mix or 10 µL PBS, were added, followed by 25 µL of the concentrations of the different mycotoxins previously prepared (plate A). Finally, 90 µL/well of exponentially growing bacteria suspension was added to each well. Then, the plates were incubated for 4h at 37°C with orbital shaking (500 rpm). After the incubation period, A₆₀₀ was measured to evaluate toxicity. Afterwards, for the determination of β-galactosidase activity, 30 µL/well of treatment plates (plate B) were transferred to new wells (plate C) containing 150 µL/well of 2-nitrophenyl-D-galactopyranoside (ONPG; Sigma-Aldrich, Switzerland) solution for the enzymatic reaction. This solution was prepared by adding 7 mL of 4.5 mg/mL ONPG solution in PBS into 28 mL of B-buffer (20.18 g/L Na_2_HPO_4_x2H_2_O, 5.5 g/L NaH_2_PO_4_xH_2_O, 0.75 g/L KCl, 0.25 g/L MgSO_4_x7H_2_O (Sigma-Aldrich, USA), 1 g/L SDS (Sigma-Aldrich, USA), 2.7 μL/mL β-mercaptoethanol (Sigma-Aldrich, USA), in distilled water). Later, 30 μL from each well of the plates B were transferred into the plates C. Plates C were incubated for 30 min at 28°C with orbital shaking (500 rpm) in the dark. After the incubation period, the reaction was stopped by adding 120 µL/well of Na₂CO₃ (1 M) and finally, A₄₂₀ was measured. Three (AOH, AME, ATX-I) or four (ALT, TeA, TEN) independent experiments were carried out per mycotoxin.

#### Data analysis

Toxicity was calculated as follows:

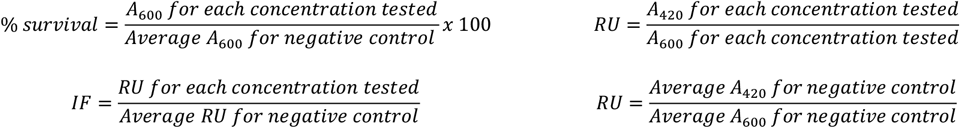

The presence of a dose-response and IF values ≥ 2 at any of the concentrations were considered as positive when the bacterial survival is above 80%. IF values < 1.5 were considered negative. IF among 1.5 and 2 were considered equivocal.

### 2.4. *In Vitro* Mammalian Cell Micronucleus Test in TK6 cells

#### Cell culture

The thymidine kinase 6 (TK6) lymphoblastoid cells (derivative from the WIL-2 lymphoblast cell line) purchased from Cytion (Heidelberg, Germany) and the European Collection of Authenticated Cell Cultures (ECACC; Salisbury, United Kingdom), were used during linear growth (exponential phase). The cells were cultured in RPMI-1640 medium supplemented with 10% heat-inactivated fetal bovine serum, 1% gentamycin, 1% glutamax, 1% sodium pyruvate, 1% non-essential amino acids, and 0.1% amphotericin B. The cell medium and all the medium supplements were purchased from Thermo Fisher Scientific (Merelbeke, Belgium) and stored in the conditions recommended by the provider.

#### Cytokinesis-Block Micronucleus (CBMN) assay in TK6 cells

The *in vitro* MN assays were performed on TK6 cells according to the OECD TG 487 (OECD 2023). Cytochalasin B (Cyt-B; Merck Life Science, Hoeilaart, Belgium) was used as a cytokinesis blocker and added to the cells after the 24h or 3h exposure period. The protocol used is similar to that described by Sanders et al. (2022) and Sanders et al. (2025). In practice, TK6 cells were seeded in 6-well plates at 20,000 cells/mL for a 24h exposure test and at 75,000 cells/mL for a 3h exposure test and incubated (37°C, 5% CO_2_). The next day, cells were exposed to increasing concentrations of the *Alternaria* toxins (1 replicate/concentration/experiment), with a maximum of 0.5% and 1% of DMSO in the medium with and without metabolic activation, respectively. For the tests with metabolic activation, a S9-mix was added to each well at a final concentration of 10% v/v. The S9-mix was prepared by dissolving the lyophilized phenobarbital/5,6-benzoflavone-induced Sprague-Dawley rat liver S9 (Trinova Biochem, Giessen, Germany) in reverse osmosis water and mixing it with the NADPH cofactor regenerating system (Regensys A and B, Trinova Biochem, Giessen, Germany) (1:9 v/v) according to the provider’s recommendations. Methyl methanesulfonate (MMS; Thermo Fisher Scientific, Merelbeke, Belgium) 36.3 µM (2 µg/mL), ethyl methanesulfonate (EMS; Merck Life Science, Hoeilaart, Belgium) 5 mM (620.77 mg/L), and aflatoxin B1 (AFB1; Merck Life Science, Hoeilaart, Belgium) 0.15 µM (4.68 ng/mL) were used as positive controls for the assays without metabolic activation for 24h and 3h, and with metabolic activation, respectively. Medium containing 0.5% or 1% DMSO was used as the negative control. After incubation for the adequate exposure time (i.e., 24h or 3h), cells were inspected visually under an inverted optical microscope (Zeis Primovert equipped with a camera axiocam 105 color) to check their aspect and the presence of precipitates before being cleared of the exposed mycotoxins by centrifugating the samples (5 min at 102*x*g) and removing the supernatant. Cells were directly re-suspended in medium for the tests without metabolic activation. For the tests with metabolic activation, cells were washed twice with 5 mL of medium (supernatant removed after centrifugation for 5 min at 102*x*g) before resuspending them in fresh medium. Cells were then exposed to 6 µg/mL of Cyt-B for 21h. Finally, the fixation of the cells was performed by adding drop-by-drop 1 mL of a pre-warmed hypotonic solution (0.075 M KCl at 35°C) for 10 min, followed by 1 mL of an ice-cold methanol-acetic acid solution (3:1 v/v) (fixator 1). After 10 min, the tubes were centrifuged for 5 min at 102*x*g. The supernatant was aspirated and the pellet was gently homogenized before adding 4 mL of fixator 1. The tubes were then centrifuged again for 5 min at 102 *x*g. The previous step was repeated with an ice-cold methanol-acetic acid mixture in a 9:1 v/v ratio (fixator 2). The supernatant was not completely removed, leaving approximately 1 mL in which the cells were resuspended before being spread to clean microscope slides (Epredia, VWR, Leuven, Belgium). Slides were left to dry overnight at room temperature before being stained with DAPI-containing Vectashield antifade mounting medium (Vector Laboratories, Labconsult, Brussels, Belgium) and analyzed using the Metafer fluorescence microscope system, controlled by Metafer4 software. At least two microscope slides (and a maximum of six) per condition were stained. Micronuclei (MN) were scored, when possible, in 5,000 binucleated cells (BNC) per slide (10,000 BNC in total per condition) except for the positive control for which minimum 2,500 BNC were checked per slide (when possible) for the presence of MN (5,000 BNC in total). The % of BNC with MN was used for statistical analysis. In addition, the cytotoxicity was assessed by determining the cytokinesis-block proliferation index (CBPI) and the percentage of relative survival. Therefore, one slide per condition was stained with 100 µL of propidium iodide at 1.5 µg/mL in phosphate buffer saline (PBS), and covered with a glass-coverslip (Gibco, Thermo Fisher Scientific, Merelbeke, Belgium) for 10 min before being rinsed twice with reverse osmosis water. This staining was performed as much as possible in the dark. A minimum of 500 cells was scored to calculate the CBPI, and the relative survival percentage was calculated by comparing the CBPI obtained at one exposure condition and the mean CBPI obtained for the negative controls, which was considered 100% survival.

#### Data analysis

The Fisher’s exact test was applied to compare the frequency of MNBNC of the mycotoxin-exposed cells to the negative control (cells exposed to solvent control) using GraphPad Prism (version 9.4.1). A p-value lower than 0.05 was assumed to be statistically significant. When at least one concentration exhibited a significant increase, the linear Chi-square trend test was applied with RStudio (version 2024.12.1 +563) to evaluate whether the increase is dose-related or not. For the R analyses, the following packages were employed: readxl (Wickham and Bryan 2025), purr (Wickham and Henry 2025), ggplot2 (Wickham 2016), dplyr (Wickham et al. 2023a), scale (Wickham et al. 2025), ggpubr (Kassambara 2025), tibble (Müller and Wickham 2025). Finally, the positive results were checked to be outside of the distribution of the historical negative control data. When all these criteria are met, the mycotoxin tested was considered positive (i.e., able to induce chromosome breaks) according to the OECD TG 487. On the contrary, if one of these criteria is not met, the mycotoxin tested was considered negative. As specified in the OECD guidelines, there is no requirement to verify a clear positive or negative result. In addition, all negative and positive controls data were evaluated for inclusion in our historical databases. Each value should ideally fall within the 95% distribution of historical data and, if not, not be an extreme outlier.

#### Fluorescence *in situ* hybridization (FISH)

A FISH protocol using pan-centromere probes labelled by FAM (fluorescein amidite) fluorescent dye (Eurogentec, Seraing, Belgium) was applied on slides obtained following the *in vitro* MN assay procedure described before. During a pre-treatment phase, slides were washed by sequential immersions in ethanol series (70%, 90% and 100%) for 5 min each and left to dry overnight. The next day, slides were successively washed in PBS (5 min), in 4% paraformaldehyde (PFA; Thermo Fisher Scientific, Merelbeke, Belgium) for 2 min, and again in PBS (3 x 5 min). The cells were then exposed to pepsin (100 µL/slide at 0.1 mg/mL) for 7 min and rinsed successively with PBS (5 min), PFA 4% (2 min), and PBS (5 min). Slides were exposed for 30 min to a 2x saline-sodium citrate (SSC) buffer (0.3 M sodium chloride and 0.03 M sodium citrate) containing 1% Triton X pre-heated at 37°C before being rinsed twice with PBS (2 x 5 min). Before the hybridization, slides were washed and dehydrated in a graded ethanol series (70%, 90%, and 100%) for 5 min each. The hybridization step was induced by adding the centromere probe (20 µL at 0.05 nmol) to each slide. Afterwards, the slides were covered with a glass coverslip and placed in a pre-heated heating block at 80°C for 3 min. Slides were then placed in a dark and humid chamber in the incubator (37°C, 5% CO_2_) for 2h. After the hybridization, the cover glass was removed, and slides were washed 2 times for 15 min in a 70% formamide/10 mM Tris (pH 7.2) solution and 3 times for 5 min in a solution made of 50 mM Tris (pH 7.2)/150 mM NaCl (pH 7.5)/0.05% Tween 20. Finally, the slides were stained after a brief wash in PBS with a DAPI mounting solution and analyzed using a fluorescent microscope (Metafer system).

Each FISH staining and analysis included negative control (cells exposed to 1% DMSO) in duplicate, two positive controls: colchicine (COLC; Merck Life Science, Hoeilaart, Belgium) at 0.015 µM (6 ng/mL) as the aneugen positive control and MMS at 36.3 µM (2 µg/mL) as the clastogen positive control, and one slide for each of four increasing concentrations of the test mycotoxin showing an effect in the *in vitro* MN assay after 24h exposure without S9. Practically, the BNCs with MN were first selected using the DAPI filter of the Metafer system. Then, the MNs were checked for the presence of a centromere (MN CENT+) or not (MN CENT-) using the FITC filter. The FISH test was repeated twice for each mycotoxin, and a maximum of 100 BNC with MN were scored per slide, when possible. The proportion of MN CENT+ and MN CENT-was compared to COLC and MMS values.

### 2.5. *In vitro* micronucleus assay in HepG2 cells

#### Cell culture

HepG2 cells (ATCC® HB-8065 ™) were obtained from the American Type Culture Collection (Manassas, VA, USA) and exhibit a doubling time of approximately 48h. Cells were cultured in DMEM F-12, supplemented with 15% inactivated fetal bovine serum, 1% penicillin/streptomycin (10.000 U/mL), 1% amphotericin B (250 µg/mL) and 2.5% HEPES buffer (all from Thermo Fisher Scientific, Waltham, MA, USA). Cultures were maintained in a humidified incubator at 37°C with 5% CO₂ and were subcultured every 2 to 3 days, when cell confluence reached approximately 70%.

#### Preliminary cytotoxicity testing

The 3-(4,5-dimethylthiazol-2-yl)-2,5-diphenyltetrazolium bromide (MTT) assay was carried out according to Mosmann (1983) with modifications. Briefly, HepG2 cells were seeded into 96-well plates at a concentration of 1×10⁵ cells/well, incubated for 24h and then exposed for 48h to a concentration range of 1, 10, 25, 50, 75, 100, and 200 µM of mycotoxin, with the exception of ATX-I in which the highest concentration was not tested due to mycotoxin precipitation. Controls included non-exposed cells (in culture medium without and with DMSO at the highest concentration used – AOH 0.67%; AME 1%; ALT and TEN 2%; ATX-I 0.5%; TeA 1%), and a positive control corresponding to cells exposed to 0.1% SDS for 1h. At 48h, cells were washed twice with PBS and 100 µL of 0.5 mg/mL MTT solution was added to each well. Following 3h incubation at 37°C in 5% CO₂, the MTT solution was removed, 100 µL of DMSO was added and, after 20 min incubation in an orbital shaker in the dark, the absorbance was measured at 570 nm with 690 nm as reference filter using a microplate reader (SpectraMax iD3, Molecular Devices). Cell viability (%) was calculated as the division of the absorbance of the sample by the absorbance of the solvent control multiplied by 100. Three independent experiments were performed, each with 6 technical replicates.

#### CBMN assay in HepG2 cells

HepG2 cells were seeded at a density of 2×10⁵ cells per well in 6-well plates, with two technical replicates for each experimental condition. Plates were incubated for 24h at 37°C and 5% CO₂ and then cells were exposed to the selected concentration range of mycotoxins in the presence of Cyt-B (2 µg/mL) for 48h at 37°C and 5% CO₂. Positive controls were treated with Vinblastine (VBL; 0.05 ng/mL), and negative controls included non-exposed cells (cells in culture medium, with and without DMSO at the highest concentration tested – AOH 0.45%; AME 0.15%; ATX-I 0.125%; ALT and TEN 1%; TeA 0.09%). At the end of the exposure, wells were washed, trypsinized, transferred to 15 mL centrifuge tube,s and centrifuged at 1200 rpm for 5 min. Hypotonic treatment was performed in the resuspended pellet by adding 5 mL of pre-warmed 0.1 M KCl drop-by-drop under vortex agitation. Then, 1 mL of a cold methanol-acetic acid (3:1) solution was added before centrifugation at 1200 rpm for 5 min, cells were resuspended, 5 mL of the same cold methanol-acetic acid solution (3:1) was added drop-by-drop under vortexing, and centrifuged again at 1200 rpm for 5 min. Finally, a second fixation step was performed using 5 mL of cold methanol-acetic acid (97:3), followed by centrifugation. Fixed cells were spread onto microscope slides and stained in 4% Giemsa solution. Slides were coded and blind scored under a bright field microscope (Axioskop 2 Plus, Zeiss, Germany) for the presence of MN, using the criteria described by Fenech (2007). At least 2000 BNC from two independent cultures were scored per treatment condition. The frequency of micronucleated binucleated cells per 1000 cells (MNBNC/1000 BNC) was determined, as well as the proportion of mono-(MC), bi-(BC) or multinucleated-cells (MTC), in order to calculate the CBPI and cytotoxicity, according to the OECD TG 487 (OECD 2023).

#### Data analysis

Statistical analyses were performed using IBM SPSS Statistics (Version 27; SPSS Inc., Chicago, IL, USA) and R (version 4.3.1 in RStudio 2023.09.1+494). For the R analyses, the following packages were employed: ggplot2 (Wickham 2016), dplyr (Wickham et al. 2023a), drc (Ritz et al. 2015), gridExtra (Baptiste and Anton 2017), grid (Team 2023), tidyverse (Wickham et al. 2019), and tidyr (Wickham et al. 2023b).

Differences among groups were assessed using one-way analysis of variance (ANOVA). When homogeneity of variance or normality were not satisfied, the Kruskal–Wallis test was applied. For dose-response analyses, both linear and log-logistic regression models were fitted, and model performance was compared using goodness-of-fit statistics (R^2^). For the CBMN assay, comparisons of the frequency of MNBNCs between mycotoxin-exposed groups and their corresponding controls were conducted using Fisher’s exact test. P-values of 0.05 or lower were considered as statistically significant.

### 2.6. γH2AX assay

#### Cell culture

Undifferentiated HepaRG cells (Biopredic International, Saint Grégoire, France) were plated in 96-well plates at a density of 9,000 cells/well and cultured according to previously established protocols (Gripon et al. 2002; Luckert et al. 2018). Briefly, the cells were cultivated in William’s E medium (Pan-Biotech, Aidenbach, Germany) supplemented with 10% fetal calf serum (Good Forte (Lot: P131102), Pan-Biotech), 100 U/mL penicillin, and 100 μg/mL streptomycin (both from Capricorn Scientific, Ebsdorfergrund, Germany), 0.05% human insulin (Pan-Biotech), and 50 μM hydrocortisone hemisuccinate (Sigma-Aldrich, Taufkirchen, Germany) at 37°C in a humidified atmosphere for 14 days. The initiation of cellular differentiation was carried out by adding 1% (v/v) DMSO to the medium for 2 days, followed by an addition of 1.7% (v/v) DMSO to the medium for up to day 28. The medium was changed every 2 to 3 days.

#### Cell viability

Cellular viability was determined using the MTT assay as previously described (Behr et al. 2020; Dietrich et al. 2019). HepaRG cells were differentiated in 96-well plates and treated with the *Alternaria* toxins (in medium containing 1.7% DMSO) for 24h. The assay was conducted in quadruplicates with at least three individual experiments. Untreated cells served as control, and medium containing 0.01% Triton X-100 served as positive control.

#### γH2AX assay

Differentiated HepaRG cells were exposed 24h to various concentrations of *Alternaria* toxins in medium containing 1.7% DMSO. Doxorubicin (DOX) (1 µM, Cayman Chemicals, Ann Arbor, MI, USA) served as a positive control and untreated cells as a negative control. Following the 24h-treatment, cells were gently washed with cold PBS and fixed with 50 μL of ice-cold methanol per well for 30 min. After fixation, the cells were washed with PBS-T (0.1% Tween-20 in PBS) and blocked with 50 μL/well of the blocking solution (1% bovine serum albumin in PBS-T) for 1h at room temperature. For antibody staining, the blocking solution was removed, and 40 μL/well of the primary antibody anti-phospho histone H2A.X (S139) (EMD Milllipore, Burlington, MA, USA), diluted 1:500 in blocking solution, was added. After 1h of incubation, the primary antibody was removed, and the wells were washed three times with PBS-T. The secondary antibody, AlexaFluor 647 (Life Technologies, Carlsbad, CA, USA), was diluted 1:400 in blocking solution, and 40 μL was added per well for another 1h incubation. The cells were then washed three times with PBS-T, followed by the addition of 50 μL DAPI (3 μM; TCI Deutschland, Eschborn, Germany) per well. After a 30-min incubation, fluorescence intensity was measured using the Celldiscoverer 7 microscope by scanning the well automatically (four pictures per well) at 10x magnification in at least three individual experiments with four replicate wells per condition. Image analysis was performed with ZEN 3.1 software. A threshold, which was equal for the analysis of each image, was set to exclude unspecific fluorescence signals.

#### Data analysis

For γH2AX, results were considered positive when there was a statistically significant two-fold induction in more than two adjacent concentrations compared to the control. If only one concentration showed a fold-induction of ≥ 2, results were considered as equivocal. For determination of differences between control and treatment groups, one-way ANOVA was performed with Dunnett’s post hoc test using GraphPad prism 10.1.2. Data were considered significantly different when *p < 0.05, **p < 0.01 or ****p < 0.0001.

## 3. Results

### 3.1. Gene mutation assay (Ames test)

The tested toxins were fully soluble in sterile DMSO, forming a homogeneous solution at the selected concentrations specific for each toxin, with no visible precipitate. No precipitation of the toxins was observed in TOP agar or on the plates. Spontaneous reversion rates (negative control) and the numbers of revertant colony induced by the reference materials, both in the absence and presence of S9 metabolic activation, were within the acceptable historical control range of the laboratory for all tested bacterial strains. No signs of contamination were detected in the toxins (at the highest concentration of each toxin), growth media, S9 mix, phosphate buffer, or agar plates.

Toxicity, indicated by a moderately reduced bacterial background or an IF below 0.5, was observed only in bacteria treated with AOH, AME, and ALT. For AOH, toxic effects were observed in strain TA97a without S9 metabolic activation, and in strains TA102 and TA1535 at the highest tested concentration (0.1 mg/plate), irrespective of metabolic activation. AME and ALT exhibited toxicity only in strain TA102 without S9 metabolic activation at their highest tested concentrations, 0.272 mg/plate and 0.292 mg/plate, respectively (Table 2).

**Table 2:** Mutagenic activity of *Alternaria* toxins in S.Typhimurium in the absence (−S9) or presence (+S9) of an external metabolizing enzyme system (10% rat liver S9 mix). Data are presented as induction factor (IF) compared to solvent control.

| Treatment<br>[mg/plate] | *Salmonella typhimurium* strain |  |  |  |  |  |  |  |  |  |
| --- | --- | --- | --- | --- | --- | --- | --- | --- | --- | --- |
|  | TA97a |  | TA98 |  | TA100 |  | TA102 |  | TA1535 |  |
|  | -S9 | +S9 | -S9 | +S9 | -S9 | +S9 | -S9 | +S9 | -S9 | +S9 |
| *AOH* |  |  |  |  |  |  |  |  |  |  |
| 0.001 | 1.76 | 0.97 | 1.08 | 0.92 | 0.85 | 0.90 | 0.86 | 1.02 | 0.67 | 1.23 |
| 0.00317 | **2.26** | 1.30 | 0.99 | 0.91 | 0.97 | 0.95 | 1.03 | 1.08 | 0.85 | 1.35 |
| 0.01001 | **3.44** | 1.92 | 1.16 | 1.13 | 0.93 | 0.97 | 0.95 | 1.14 | 1.67 | 1.54 |
| 0.03165 | **4.64** | **3.11** | 1.38 | 1.05 | 1.23 | 1.27 | 1.61 | **2.02** | 1.44 | 1.12 |
| 0.1 | 0.51<sup>b</sup> | **2.80** | 1.47 | 0.57<sup>a</sup> | 1.15<sup>a</sup> | 1.32<sup>a</sup> | 0.55<sup>b</sup> | 2.86<sup>b</sup> | 0.59<sup>b</sup> | 0.15<sup>b</sup> |
| PC | **3.59** | **3.90** | **7.50** | **8.95** | **3.04** | **5.39** | **2.26** | **2.54** | **18.09** | **21.05** |
| *AME* |  |  |  |  |  |  |  |  |  |  |
| 0.0027 | 1.41 | 1.46 | 1.19 | 1.04 | 1.03 | 1.05 | 1.05 | 1.16 | 0.83 | 0.84 |
| 0.0086 | **2.11** | **2.15** | 0.84 | 1.17 | 1.07 | 1.19 | 1.06 | 1.19 | 0.86 | 0.75 |
| 0.027 | **3.06** | **3.65** | 0.85 | 1.32 | 1.21 | 1.25 | 1.11 | 1.17 | 0.94 | 0.75 |
| 0.086 | **3.38** | **4.14** | 1.27 | 1.23 | 1.08 | 1.24 | 0.97 | 1.27 | 0.89 | 1.00 |
| 0.272 | **3.58** | **4.63** | 0.66 | 1.26 | 1.09 | 1.22 | 0.42 | 0.62 | 0.71 | 0.63 |
| PC | **2.54** | **2.63** | **14.84** | **4.01** | **4.27** | **5.21** | **3.59** | **2.88** | **11.71** | **7.76** |
| *ALT* |  |  |  |  |  |  |  |  |  |  |
| 0.0029 | 1.04 | 1.02 | 0.79 | 1.00 | 0.82 | 0.95 | 1.01 | 1.01 | 0.83 | 1.25 |
| 0.0092 | 1.02 | 1.16 | 0.87 | 0.93 | 0.85 | 0.87 | 0.99 | 1.07 | 0.98 | 0.75 |
| 0.029 | 1.13 | 1.08 | 1.05 | 0.98 | 0.87 | 0.85 | 0.91 | 1.06 | 0.71 | 0.83 |
| 0.092 | 1.06 | 1.04 | 0.98 | 0.79 | 1.06 | 0.85 | 0.82 | 0.84 | 0.78 | 1.22 |
| 0.292 | 0.96 | 1.07 | 1.12 | 1.14 | 0.76 | 0.98 | 0.49 | 0.65 | 0.83 | 1.06 |
| PC | **2.90** | **2.68** | **9.60** | **3.32** | **4.86** | **8.41** | **3.82** | **3.47** | **14.37** | **8.77** |
| *ATX-I* |  |  |  |  |  |  |  |  |  |  |
| 0.0001 | 1.21 | 1.01 | 1.15 | 0.88 | 1.23 | 0.84 | 1.05 | 1.06 | 1.00 | 0.90 |
| 0.0003169 | 1.15 | 0.93 | 1.08 | 0.92 | 1.10 | 1.00 | 1.04 | 0.99 | 1.21 | 0.90 |
| 0.001 | 1.23 | 0.99 | 1.05 | 1.16 | 1.03 | 0.95 | 1.00 | 1.00 | 0.86 | 1.30 |
| 0.00316 | 1.43 | 1.03 | 1.58 | 1.03 | 1.27 | 0.98 | 1.33 | 0.99 | 0.79 | 0.83 |
| 0.010 | **2.03** | 1.12 | 1.58 | 1.05 | 1.37 | 1.04 | **2.23** | 1.12 | 1.32 | 1.23 |
| PC | **3.24** | **3.55** | **15.41** | **4.96** | **3.82** | **5.91** | **2.81** | **3.42** | **17.86** | **22.73** |
| *TeA* |  |  |  |  |  |  |  |  |  |  |
| 0.025 | NA | NA | 1.08 | 1.17 | 1.08 | 0.96 | NA | NA | NA | NA |
| 0.079 | NA | NA | 1.11 | 1.08 | 0.98 | 1.21 | NA | NA | NA | NA |
| 0.25 | NA | NA | 0.98 | 1.21 | 0.94 | 1.16 | NA | NA | NA | NA |
| 0.791 | NA | NA | 0.98 | 1.07 | 1.12 | 0.92 | NA | NA | NA | NA |
| 2.5 | NA | NA | 1.10 | 0.97 | 1.10 | 1.08 | NA | NA | NA | NA |
|  | NA | NA | 8.32 | 4.58 | 3.54 | 7.36 | NA | NA | NA | NA |
| *TEN* |  |  |  |  |  |  |  |  |  |  |
| 0.0041 | 0.98 | 0.97 | 0.87 | 1.08 | 0.95 | 0.86 | 0.92 | 0.98 | 0.80 | 1.16 |
| 0.0129 | 1.01 | 1.07 | 1.11 | 1.24 | 0.95 | 0.82 | 0.87 | 1.06 | 0.80 | 1.21 |
| 0.041 | 1.08 | 0.98 | 0.74 | 0.93 | 0.90 | 0.94 | 0.95 | 1.11 | 0.63 | 1.08 |
| 0.131 | 1.07 | 1.02 | 0.92 | 1.09 | 0.89 | 0.92 | 0.75 | 1.02 | 0.78 | 0.97 |
| 0.415 | 1.05 | 0.99 | 0.81 | 0.91 | 0.97 | 0.99 | 0.54 | 0.85 | 0.83 | 0.97 |
| PC | **2.87** | **2.68** | **9.04** | **3.32** | **2.86** | **8.41** | **3.44** | **3.71** | **16.70** | **8.70** |

Bolded values: IF $\geq 2$ (TA97a, TA98, TA100, TA102) or IF $\geq 3$ (TA1535). <sup>a</sup> slightly reduced bacterial background, <sup>b</sup> moderately reduced bacterial background, NA: not available; Positive controls (PC): without S9: TA97a and TA98 = 4-nitroquinoline-N-oxide (0.25 $\mu\text{g}/\text{plate}$ ), TA100 = sodium azide (0.25 $\mu\text{g}/\text{plate}$ ), TA1535 = sodium azide (0.125 $\mu\text{g}/\text{plate}$ ), TA102 = mitomycin C (0.5 $\mu\text{g}/\text{plate}$ ); with S9: TA97a = benzo(a)pyrene (5 $\mu\text{g}/\text{plate}$ ), TA98 and TA100 = benzo(a)pyrene (2.5 $\mu\text{g}/\text{plate}$ ), TA102 = 2-aminoanthracene (10 $\mu\text{g}/\text{plate}$ ), TA1535 = 2-aminoanthracene (5 $\mu\text{g}/\text{plate}$ )

Table 2 presents the IF compared to the solvent control. The number of revertants including mean and SD can be found in Table S1 in the supplementary material. AOH and AME induced a > 2-fold increase in the mean number of revertants in *S.Typhimurium* strain TA97a, both in the absence and presence of metabolic activation system. In addition, AOH also induced a > 2-fold increase in strain TA102 in the absence of metabolic activation system (at 0.03165 mg/plate). Similarly, ATX1 at 0.01 mg/plate induced a > 2-fold increase in *S.Typhimurium* strains TA97a and TA102 in the absence of metabolic activation system (Table 2). These results indicate that AOH and ATX1 are capable of inducing both frameshift mutations and base-pair substitutions, whereas AME appears to induce only frameshift mutations. Contrary, TEN and ALT, at concentrations up to 0.415 mg/plate and 0.292 mg/plate respectively, did not induce a > 2-fold increase in strains TA97a, TA98, TA100, or TA102, nor a > 3-fold increase in strain TA1535, compared to the solvent control, regardless of S9. Similarly, TeA up to 2.5 mg/plate did not induce a > 2-fold increase in revertant counts in strains TA98 and TA100 (Table 2).

### 3.2 SOS/umu Assay

The SOS/umu test was employed as a screening for mutagenicity, given its high concordance with the Ames test (OECD 2020; Reifferscheid and Heil 1996). Moreover, its application has been demonstrated for use with mycotoxins (Al-Ayoubi et al. 2023; Alonso-Jauregui et al. 2021; Alonso-Jauregui et al. 2022). All the controls used in all the SOS/umu experiment were correct (IF < 2 for negative and IF > 2 for positive controls) in absence of toxicity to bacteria (survival >80%). The wells exhibiting precipitation or toxicity (bacterial survival below 80%) were excluded from the analysis. Fig 2 presents the results of the SOS/umu assay of all tested *Alternaria* toxins. AOH was clearly positive with IF above 2 and a clear dose response in the absence and presence of metabolic activation. Precipitation was observed in the initial dilution (plate A) for the four highest concentrations (from 24.22 to 193.75 µg/mL). Following the bacterial incubation period, the formation of some precipitates was also noted at the fifth concentration (12.11 µg/mL). AME and ATX-I were classified as equivocal. This classification was based on their IF, which exceeded 1.5 but remained below 2 at specific concentrations, lacking a clear concentration-response. Despite the absence of precipitate during the initial dilution (plate A), AME exhibited some formation of precipitate at the three highest concentrations following bacterial incubation. ATX-I did not exhibit precipitation in the assay; however, the tested concentrations were selected based on previous experiments in which ATX-I precipitated at 22 µg/mL. The remaining *Alternaria* toxins tested (ALT, TeA, and TEN) were clearly negative, consistently demonstrating IFs below 1.5 across all tested conditions.

**Fig 2:**
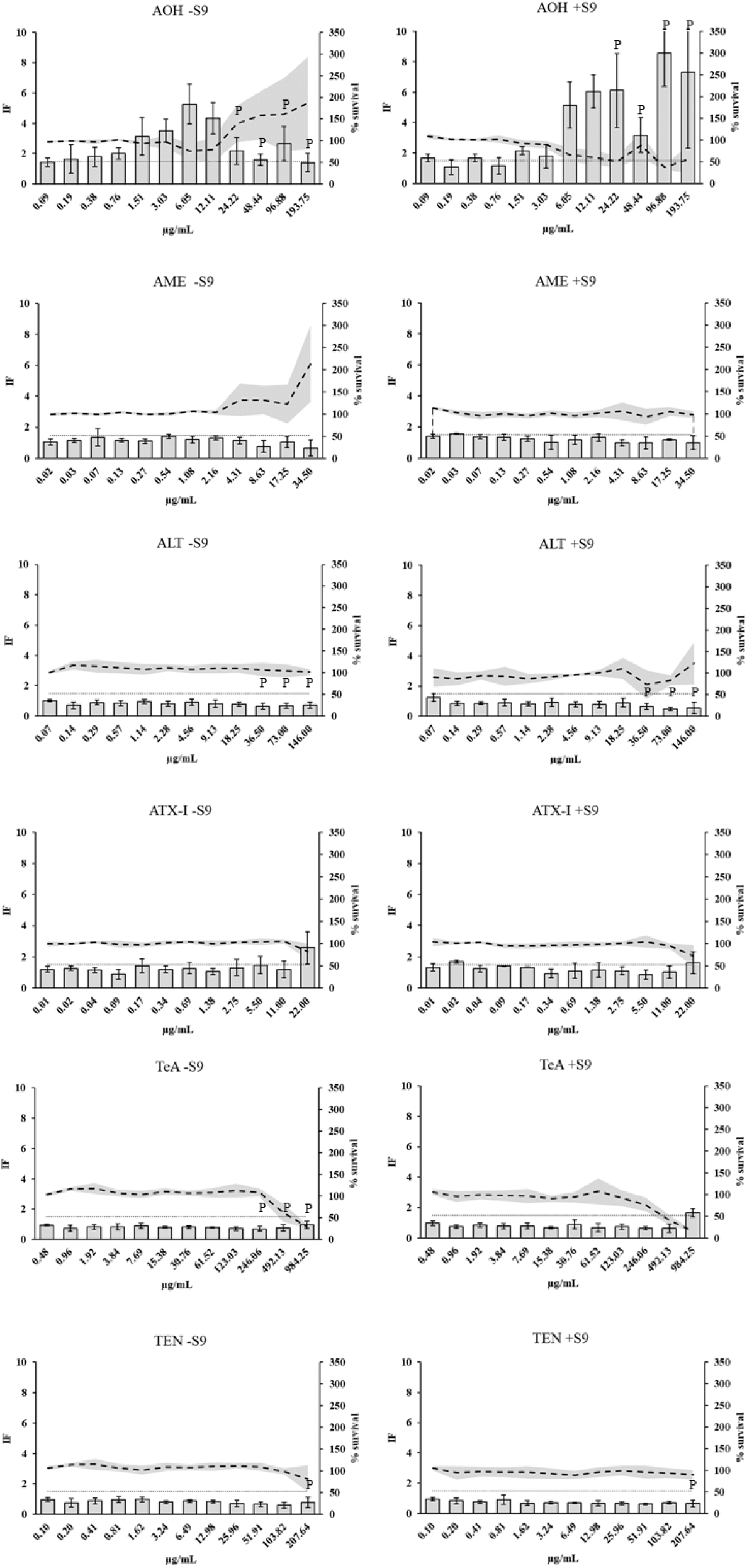
Results of SOS/umu test for *Alternaria* toxins without (-S9) or with metabolic activation (+S9). Dashed lines represent mean ± SD of bacterial survival expressed as percentage (%) (dashed line) and bars represent mean ± SD of induction factor (IF) out of three (AOH, AME, ATX-I) or four (ALT, TEN, TeA) independent experiments. Concentrations are considered non-toxic if bacterial survival is > 80%. A compound is considered genotoxic if the IF is ≥ 2 at non-toxic concentrations for the bacteria in any of the conditions tested and shows a dose-response. IF values ≥ 1.5 in absence of dose response are classified as equivocals. P: precipitates observed in the initial dilutions (plate A). Dotted line: IF = 1.5.

### 3.3. Micronucleus assay in TK6 cells

The selected *Alternaria* toxins were tested using the *in vitro* MN assays in TK6 cells under the conditions recommended in the OECD TG 487: 24h without S9 and 3h with S9. In case of negative results in both conditions, the *Alternaria* toxins were tested for 3h without S9 (OECD 2023).

#### 24h without metabolic activation

ALT, AME, AOH, ATX-I, TeA, and TEN were tested at least once without metabolic activation for 24h exposure. display the results of one experiment for each mycotoxin. AOH, AME, ATX-I, and TeA induced a statistically significant concentration-dependent increase in the percentage of BNCs with MN in TK6 cells exposed to the mycotoxin for 24h without metabolic activation compared to the negative control (Fig 3). These results indicate that all four mycotoxins induce chromosome damage *in vitro* without metabolic activation. ALT and TEN were tested up to 100 µM and 200 µM, respectively. For ALT, a significant increase in the percentage of BNCs with MN compared to the negative control was observed at 1.25 µM, while a significant decrease was observed for TEN at 100 µM. However, these changes were isolated events, and no concentration-dependent effects were observed for both compounds. Consequently, the observed effects for ALT and TEN were considered to be not biologically relevant, and both compounds were concluded to be negative in the *in vitro* MN assay up to the maximum tested concentration (Fig 3). See Table S2 to S7 in the supplementary material for more comprehensive data from the CBMN assay.

**Fig 3:**
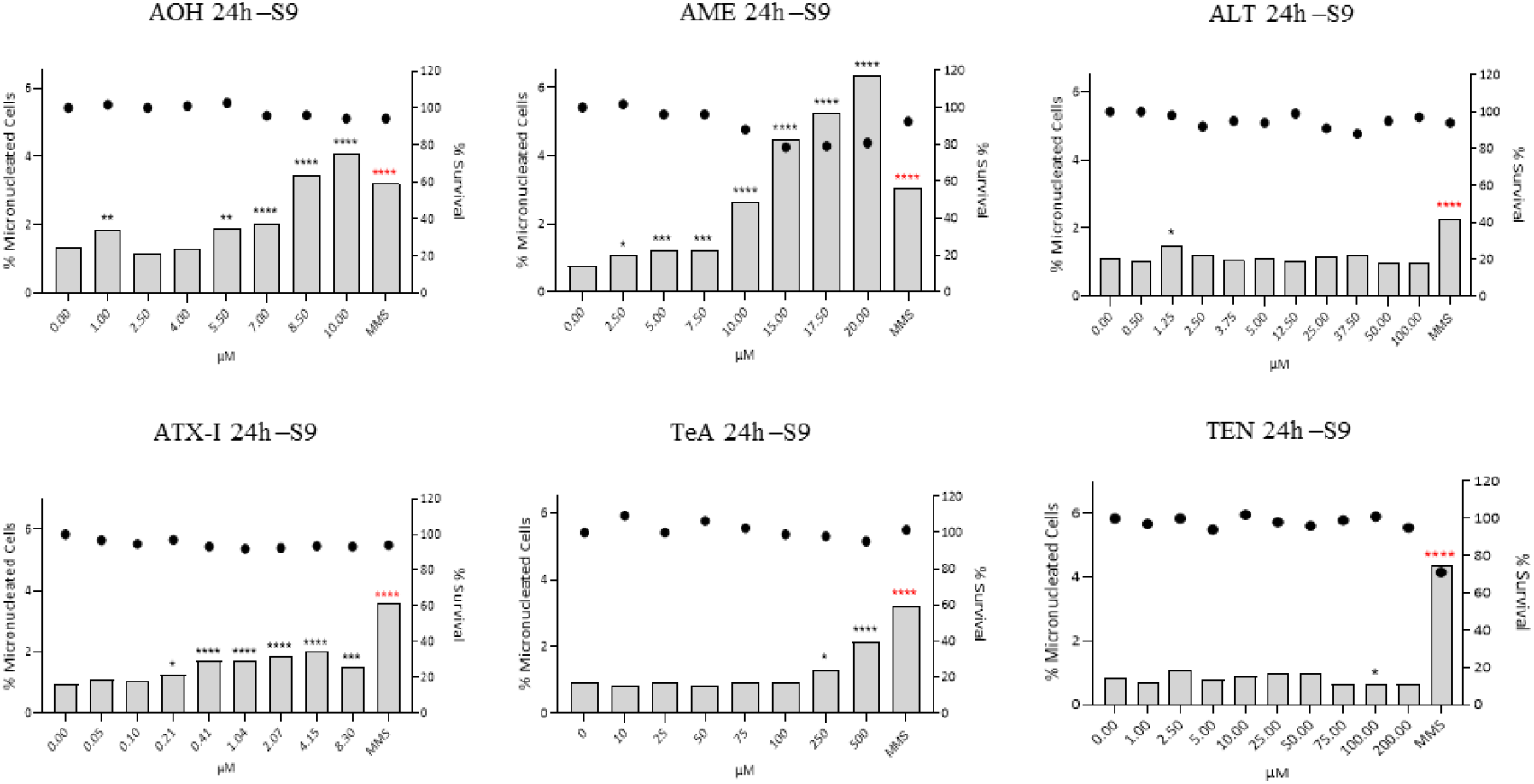
Percentage of micronucleated binucleated cells (bars) and percentage of survival calculated based on the CBPI (dots) for each condition of the individual in vitro MN assay (n=1) in TK6 cells after 24h exposure without metabolic activation (-S9). *p < 0.05, **p < 0.01, ***p < 0.001, ****p < 0.0001.

#### 3h with metabolic activation

In the *in vitro* MN assay in TK6 cells with metabolic activation (+S9), no statistically significant increase in the percentage of BNCs with MN was observed for ATX-I up to 100 µM, the maximum concentration tested in this experiment (Fig 4). As the maximum concentration induced about 60% cytotoxicity, these results suggest that ATX-I does not induce chromosome damage *in vitro* in the presence of metabolic activation (+S9). For TeA, a statistically significant increase in the percentage of BNCs with MN compared to the negative control was observed at 500 µM. However, no effect was present at concentrations of TeA above 500 µM, indicating the absence of a concentration-response effect. Also, no cytotoxicity was observed at the highest concentration tested (1000 µM). Similar results were obtained (i.e., no dose-dependent increase in the MN formation) for ALT and TEN up to 50 µM and 100 µM, respectively. The stock solution concentration did not allow for testing higher concentrations. In contrast, AOH and AME increased the percentage of BNCs with MN starting at 25 µM up to the highest concentration of 100 µM AOH or 125 µM AME (Fig 4). Notably, the effect size decreased at the highest concentration tested for AME, which may be explained by the precipitates observed under the microscope at this concentration. See Table S8 to S13 in the supplementary material for more comprehensive data from the CBMN assay.

**Fig 4:**
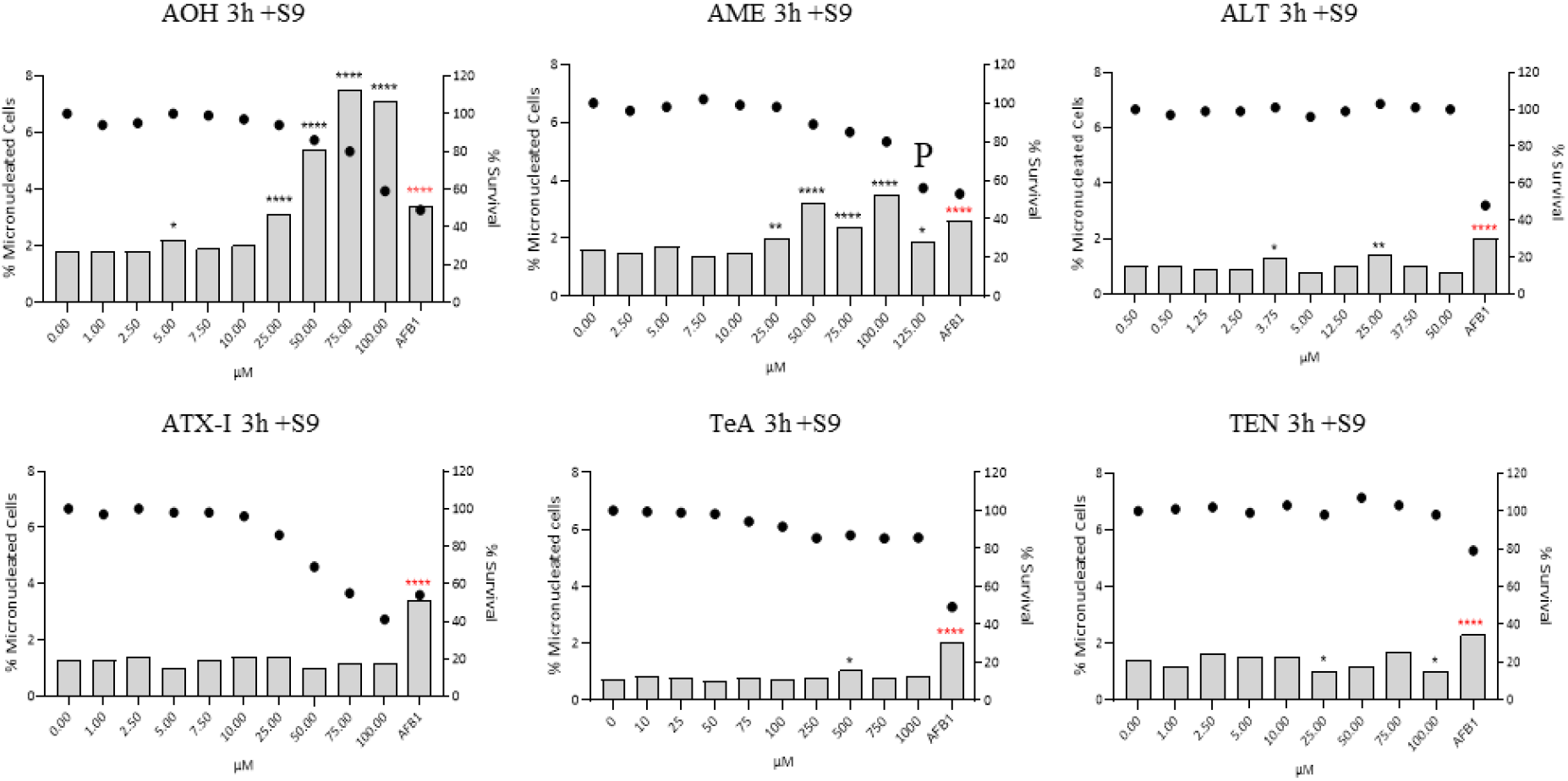
Percentage of micronucleated binucleated cells (bars) and percentage of survival calculated based on the CBPI (dots) for each condition of the individual in vitro MN assay (n=1) in TK6 cells after 3h exposure with metabolic activation (+S9). P: presence of precipitation. *p < 0.05, **p < 0.01, ****p < 0.0001.

#### 3h without metabolic activation

Since ALT and TEN were negative in both the *in vitro* MN assay with and without metabolic activation, the two mycotoxins were also tested in the 3h-exposure scenario without metabolic activation. ALT and TEN were tested up to 200 µM. Neither ALT nor TEN induced a concentration-dependent increase in MN formation in BNC, and all the %MNBNC values are included in the 95% distribution of the negative control historical data (Fig 5). No cytotoxicity was observed at the highest concentration tested. See Table S14 and Table S15 in the supplementary material for more comprehensive data.

**Fig 5:**
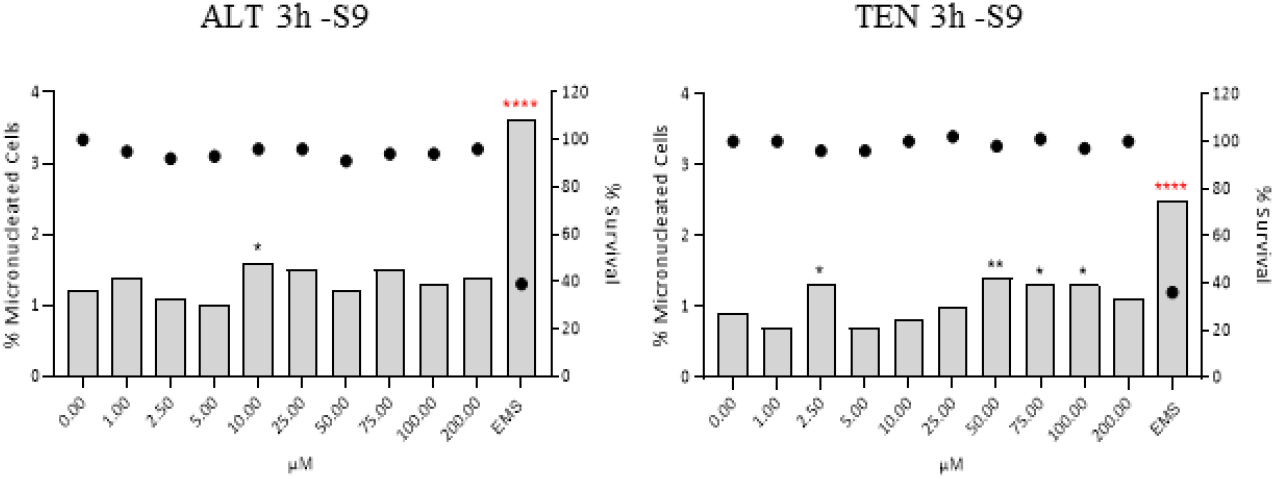
Percentage of micronucleated binucleated cells (bars) and percentage of survival calculated based on the CBPI (dots) for each condition of the individual in vitro MN assay (n=1) after 3h exposure without metabolic activation (-S9) obtained for ALT and TEN. *p < 0.05, **p < 0.01, ****p < 0.0001.

#### Fluorescence in situ hybridization (FISH)

The FISH test was conducted using four increasing concentrations of AOH, AME, ATX-I, and TeA on the slides obtained after the *in vitro* MN assay on TK6 cells after 24h exposure without metabolic activation. The selected four concentrations showed an increase in the MN formation in the *in vitro* MN assay, except for TeA, for which the increase was only observed at the two highest concentrations (Fig 3). In each FISH test, slides with TK6 cells exposed to COLC and MMS were included as aneugenic and clastogenic positive control, respectively, and cells exposed to 1% DMSO were used as a negative control. The results of the repeat FISH assays (2 slides/conditions in total) were combined for each mycotoxin (Fig 6). In all FISH tests, the proportion of MN CENT+ ranged from 64% to 68.3% for COLC, while it was between 15% and 31% for MMS. All the tested concentrations for AOH, AME, ATX-I, and TeA showed a majority of MN CENT-with a percentage of MN CENT+ between 11.6% and 33.3%. The proportion of MN CENT- and MN CENT+ obtained for each concentration of the tested mycotoxins was not significantly different from the proportion obtained for the respective MMS in the same experiment and statistically different than the proportion obtained for COLC (details on statistical analysis in supplementary material). These results suggest a clastogenic mode of action (i.e., the induction of structural chromosome aberrations), rather than aneugenicity for AOH, AME, ATX-I, and TeA at the tested concentrations. See Table S16 to S19 in the supplementary material for more comprehensive data from the FISH assay.

**Fig 6:**
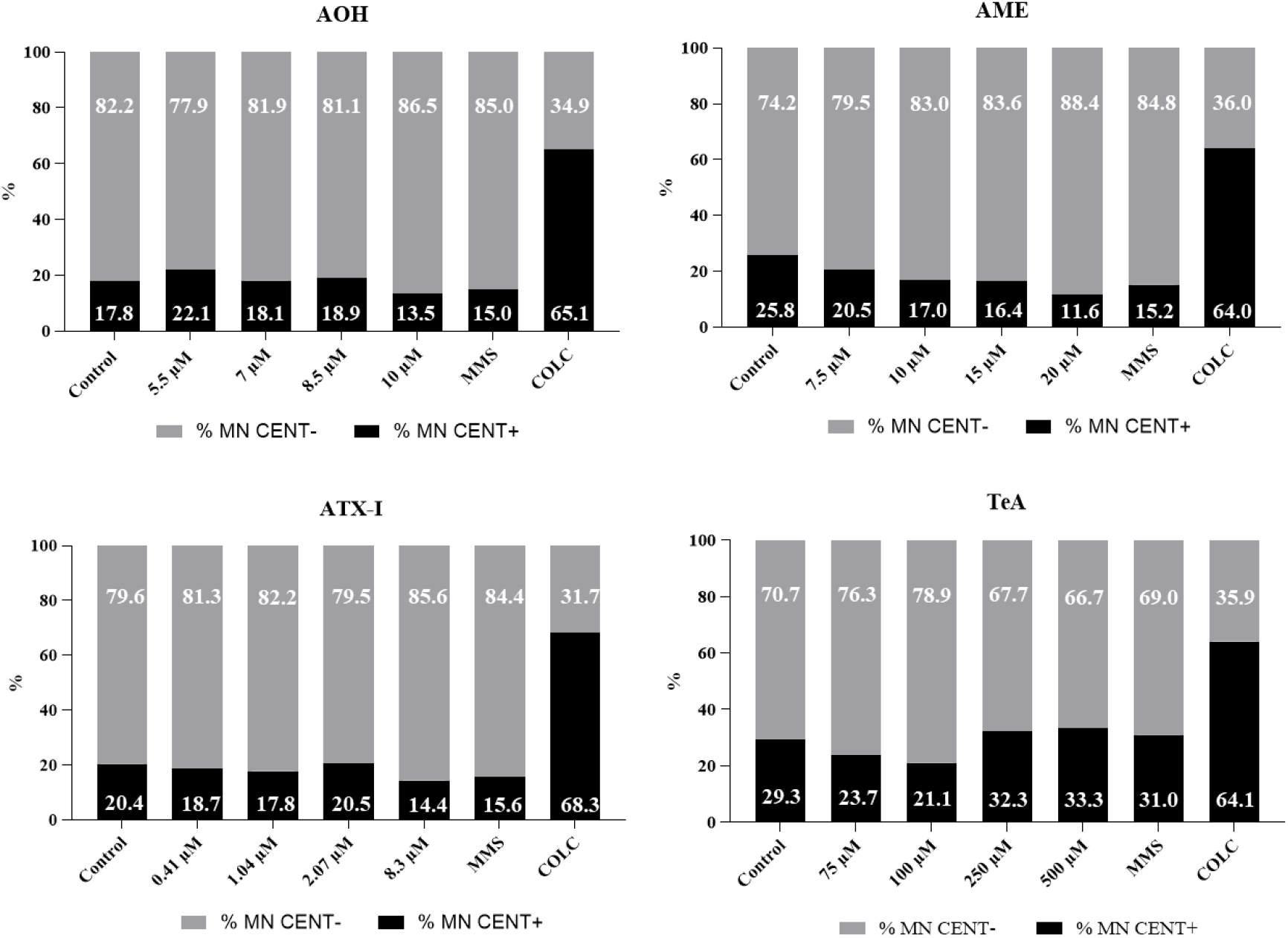
Percentage of micronuclei with (MN CENT+) and without (MN CENT-) centromere analysed after FISH staining of TK6 cells exposed for 24h without metabolic activation to different concentrations of AOH, AME, ATX-I, and TeA. For each staining, MMS and COLC were included as clastogenic and aneugenic positive controls together with a negative control consisting of cells exposed to medium with 1% DMSO.

### 3.4. Micronucleus assay in HepG2 cells

#### Preliminary cytotoxicity testing

The MTT results obtained after 48h exposure are shown in Fig S1 in the supplementary material. Regarding ALT and TEN, no statistically significant cytotoxicity was observed up to 200 µM, with a slight decrease in cell viability at this concentration (64% and 65% cell viability, respectively). AOH exhibited slight cytotoxicity at 75 µM with 59% cell viability, moderate cytotoxicity at 100 µM with 48% viability and severe cytotoxicity at 200 µM (viability 24%). AME results indicate clear cytotoxicity above 25 µM. ATX-I was more cytotoxic, with 66% cell viability at 10 µM. Higher concentrations further intensified cytotoxicity, with cell viability falling to 25% at 100 µM. TeA showed a significant decrease in the cell viability above 200 µM. At 400 µM, cell viability was below 50% of the non-exposed. Dose-response relationships were modeled using the MTT-derived viability data, obtaining an IC_50_ of 289.21 µM for ALT (extrapolated), 116.78 µM for AOH, 20.06 µM for AME, 56.84 µM for ATX-I, 446.31 µM for TeA (extrapolated) and 273.26 µM for TEN (extrapolated).

#### CBMN assay

The genotoxicity of the selected *Alternaria* toxins was investigated in HepG2 cells exposed for 48h, using the In Vitro Mammalian Cell Micronucleus Test, according to the OECD TG 487 (OECD 2023).

Prior to the assays, the solubility of each mycotoxin in culture medium was evaluated. Test solutions were prepared at 200 μM for ALT, AOH, TeA and TEN, and up to 800 μM for AME and TeA. The presence of precipitates or aggregates was assessed by naked eye evaluation and microscopically (1000X), after 24h and 48h incubation. There was no visible formation of precipitates at any of the timepoints tested for AME, AOH, ALT, TeA, and TEN. In contrast, ATX-I showed clear precipitate formation within the first 24h. Subsequently, precipitation was noted in all cultures exposed to concentrations higher than 1 μM, which did not enable the analysis of MN at those concentrations. No evidence of insolubility was observed for TEN and ALT up to 250 μM. However, the percentage of DMSO in culture (up to 1%) limited the possibility of testing concentrations above 100 μM. For AOH and TeA, the level of cytotoxicity based on the CBPI values reached 55 ± 5% cytotoxicity, as recommended in the OECD TG 487 (OECD 2023).

The results are presented in Fig 7 and comprehensive data are presented in Table S20 to S25 in the supplementary material.

**Fig 7:**
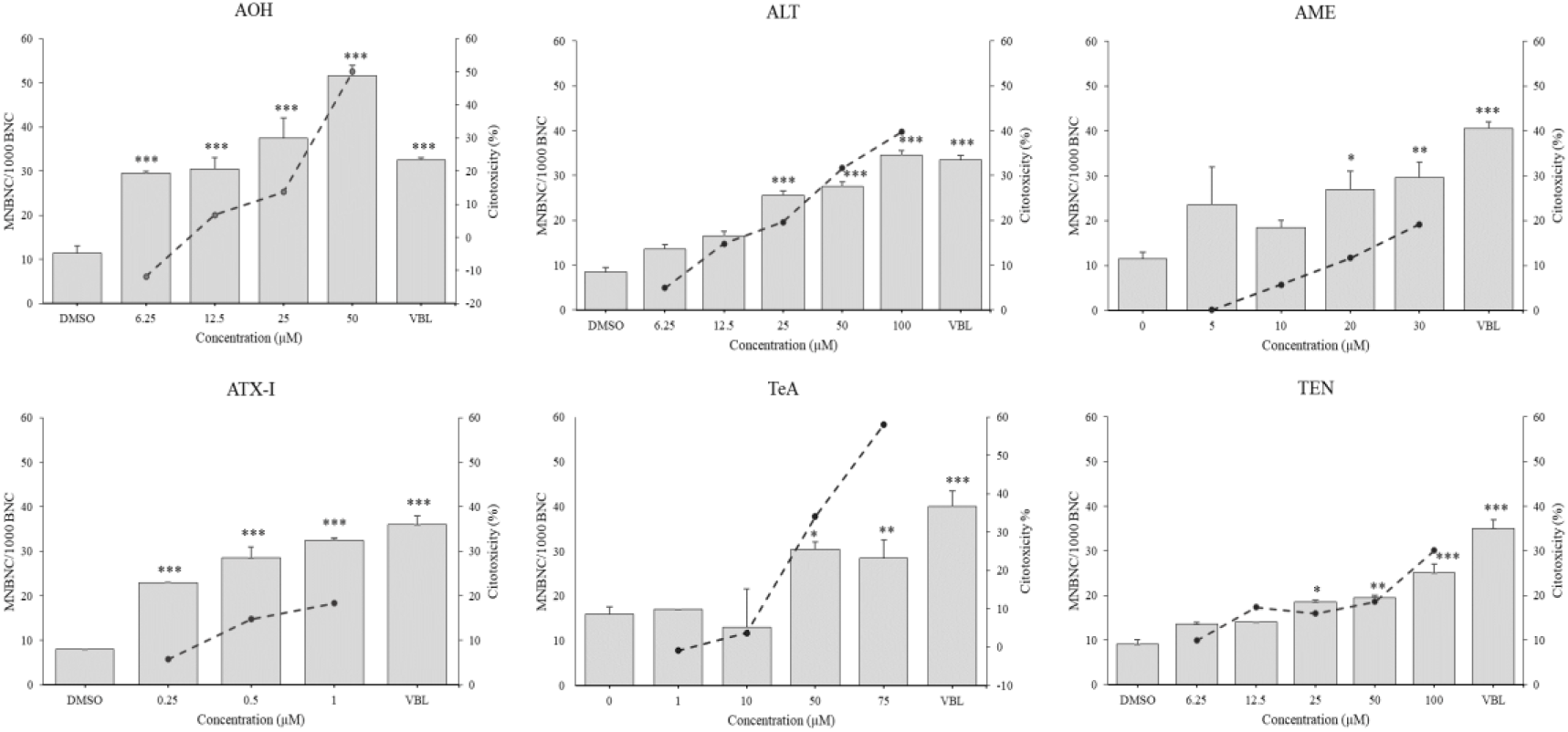
Micronucleus results and cytotoxicity after 48h-exposure to the *Alternaria* toxins, assessed by the CBMN assay. Frequencies of micronucleated binucleated (MNBNC) cells per 1000 binucleated cells (BNC) are shown on the left y-axis. The % cytotoxicity is shown on the right y-axis. Data are presented as mean + SD (n = 2 replicates). VBL: vinblastine. *p ≤ 0.05, **p ≤ 0.01, ***p ≤ 0.001.

The mean percentage of BNC in the vehicle control (DMSO) was 62.99, ranging from 52.60 ± 0.09 to 76.65 ± 0.05. ALT and TEN induced a dose-dependent increase in MNBNC frequency, which was statistically significant compared with the vehicle control, for the treatment with 25 µM and concentrations above. At the highest concentration tested, a 4-fold induction in the frequency of MNBNCs was observed for ALT. In addition, the frequency of nuclear buds was also statistically significant above 12.5 µM and 50 µM for ALT and TEN, respectively. AOH produced significant increases in the MNBNC frequency at all concentrations tested compared with the vehicle control. A significant induction of nuclear buds was also produced by 12.5 µM and concentrations above. Regarding AME, a significant increase in MNBNC was observed following cells exposure to 20 µM and 30 µM. ATX-I showed genotoxicity at all tested concentrations. The two highest concentrations tested (10 and 25 µM) could not be analyzed due to visible precipitation in the form of needle-like crystals of ATX-1 in the stained slides, which compromised exposure homogeneity. TeA was able to induce a significant increase in MNBNC in concentrations above 50 µM. A 1.8-fold increase was noted at the higher concentration.

All mycotoxins tested positive in the long-term exposure protocol, using simultaneous exposure of cells to the test substance and Cyt-B throughout the treatment period of 1.5–2 cell cycles. Therefore, according to OECD TG, no further testing upon short-term exposure protocol was needed to conclude on the capacity of each one of the toxins to induce chromosomal damage in HepG2 cells.

Atypical nuclear morphologies were also observed after AOH, ALT, ATX-I, and TEN exposures, but only reaching a clear significance in the case of ALT, right from the lowest concentration tested (6.25 μM, *p* ≤ 0.05; 12.5 μM, *p* ≤ 0.01, and 25, 50 and 100 μM, *p* ≤ 0.001). Some of these nuclear abnormalities could be classified as circular nuclei, and a residual fraction as horse-shoe shaped nuclei and fused nuclei (Fig 8).

**Fig 8:**
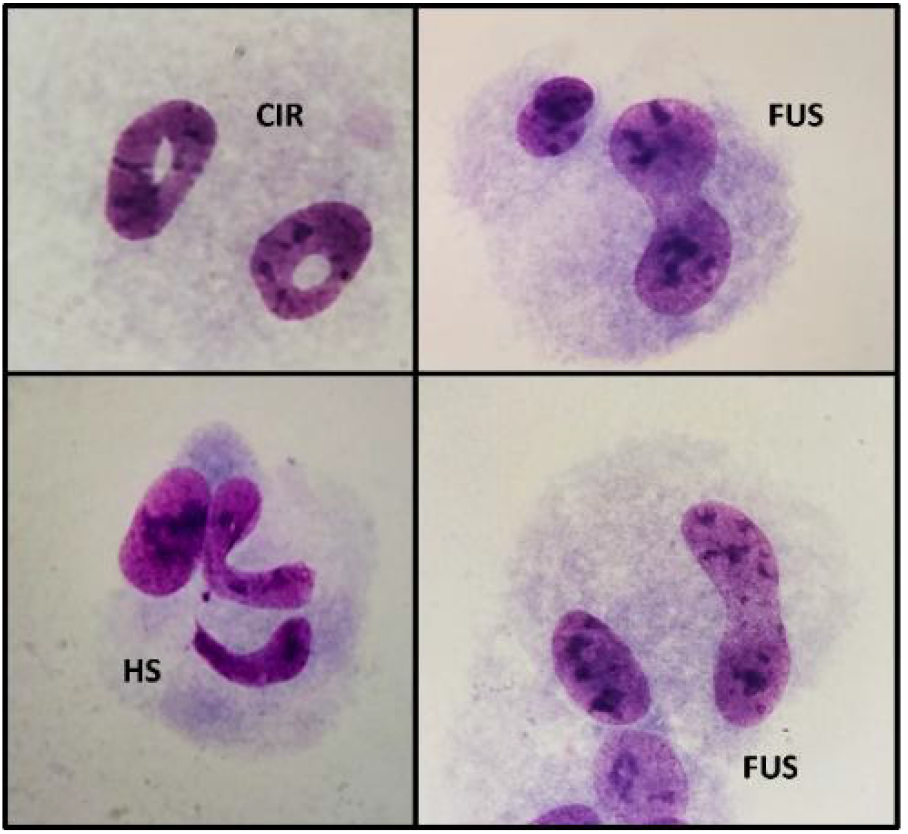
Representative Giemsa-stained cells from the CBMN assay displaying rare nuclear abnormalities. Circular nuclei (CIR), fused nuclei (FUS) and horse-shoe shaped nuclei (HS) (magnification 400X)

### 3.5. γH2AX assay

For range finding prior to the γH2AX assay, the cytotoxic potential of the selected *Alternaria* toxins was determined via MTT assay in differentiated HepaRG cells. The highest tested concentration was obtained by diluting the stock solutions (Table 1) 1:100 in the assay medium. However, ATX-I showed precipitation at 200 and 150 µM, therefore the highest test concentration was limited to 100 µM. The highest soluble concentration of each toxin was selected, followed by a series of six concentrations obtained through two-fold serial dilutions. Only AOH and ATX-I showed a reduction in metabolic activity to less than 80% compared to untreated control cells with IC_50_ value of 441.3 µM and 129.8 µM, respectively. All other toxins did not exhibit cytotoxic effects in HepaRG cells (Fig. S2).

The γH2AX assay was performed to detect primary DNA damage. The phosphorylation of the histone H2AX was quantified as an early response marker for the induction of double-strand breaks. Therefore, differentiated HepaRG cells were treated with the selected mycotoxins for 24h, followed by immunostaining and automated image aquisition and analysis. To assess the cytotoxicity in each well, DAPI staining of the cell nucleus was performed. As shown in Fig 9, AOH, ATX-I and TeA were able to induce the phosphorylation of γH2AX. However, AOH and TeA induced γH2AX phosphorylation with a ≥ 2-fold increase observed only at the highest test concentration (300 µM for AOH and 1000 µM for TeA). ATX-I induced γH2AX concentration-dependent up 3.5-fold compared to the control. AME, ALT and TEN showed no effect on the phosphorylation status of γH2AX at the tested concentrations.

**Fig 9:**
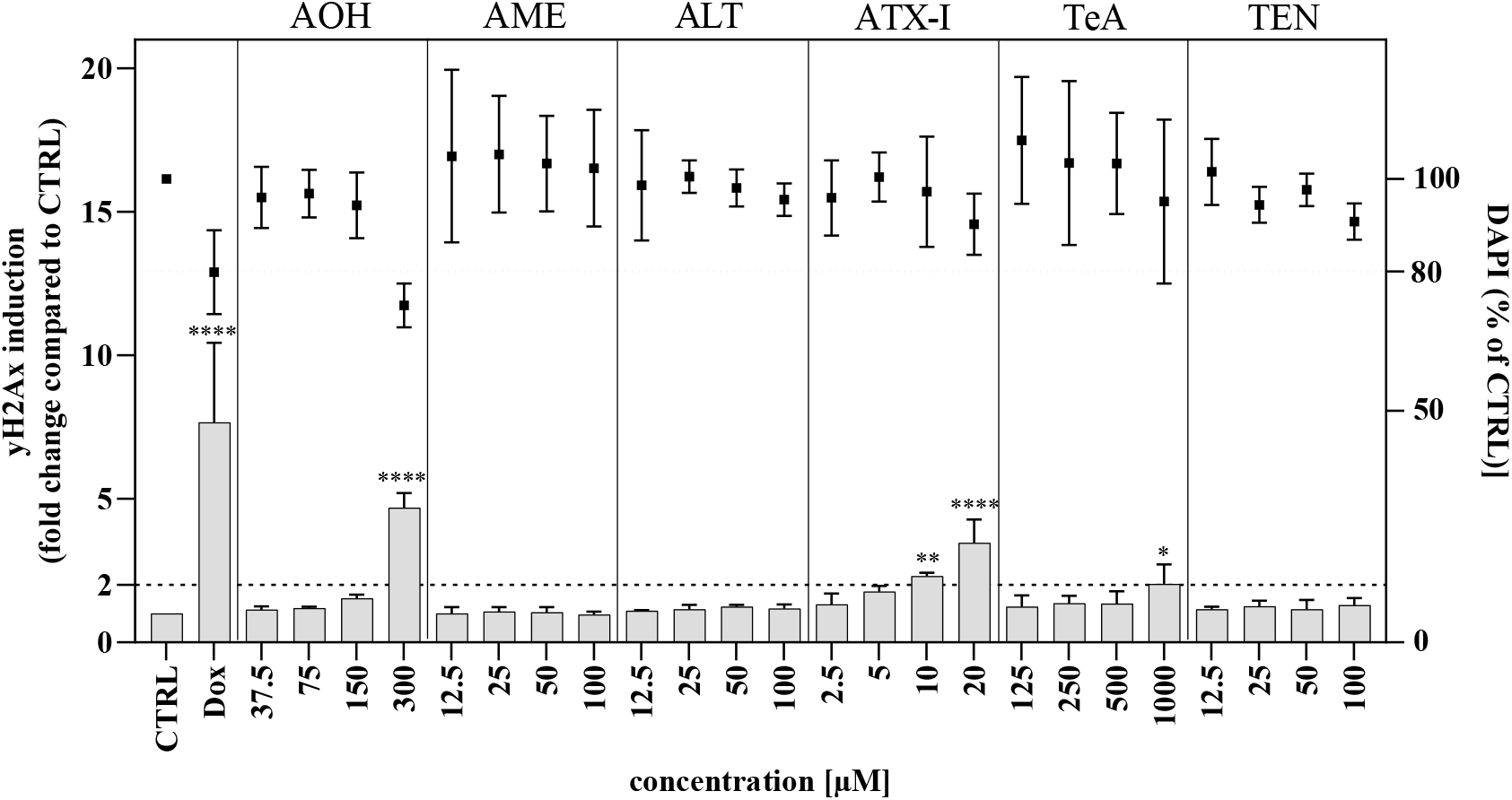
Results of the γH2AX assay in HepaRG cells. The cells were treated with *Alternaria* toxins for 24h. Bars represent the fluorescence intensity of phosphorylated histone H2AX (γH2AX) and dots represent DAPI staining normalized to untreated cells (CTRL). Cells exposed to doxorubicin (DOX, 1 µM) served as positive control. Data are presented as mean + SD (γH2AX) and mean ± SD (DAPI) with *p ≤ 0.05, **p ≤ 0.01, ****p ≤ 0.0001.

### 3.6. Overview of results

From **Fehler! Verweisquelle konnte nicht gefunden werden.**, it becomes apparent that, in the tested conditions, AOH, AME and ATX-I are genotoxic *in vitro*, revealing both mutagenic and chromosomal damage potential. ALT and TEN showed capacities to induce chromosomal damage in liver cells only, not in lymphoblastoid cells, and were not mutagenic in bacteria. TeA was also not mutagenic while it induced chromosomal damage both in liver and lymphoblastoid cells.

**Table 2:**
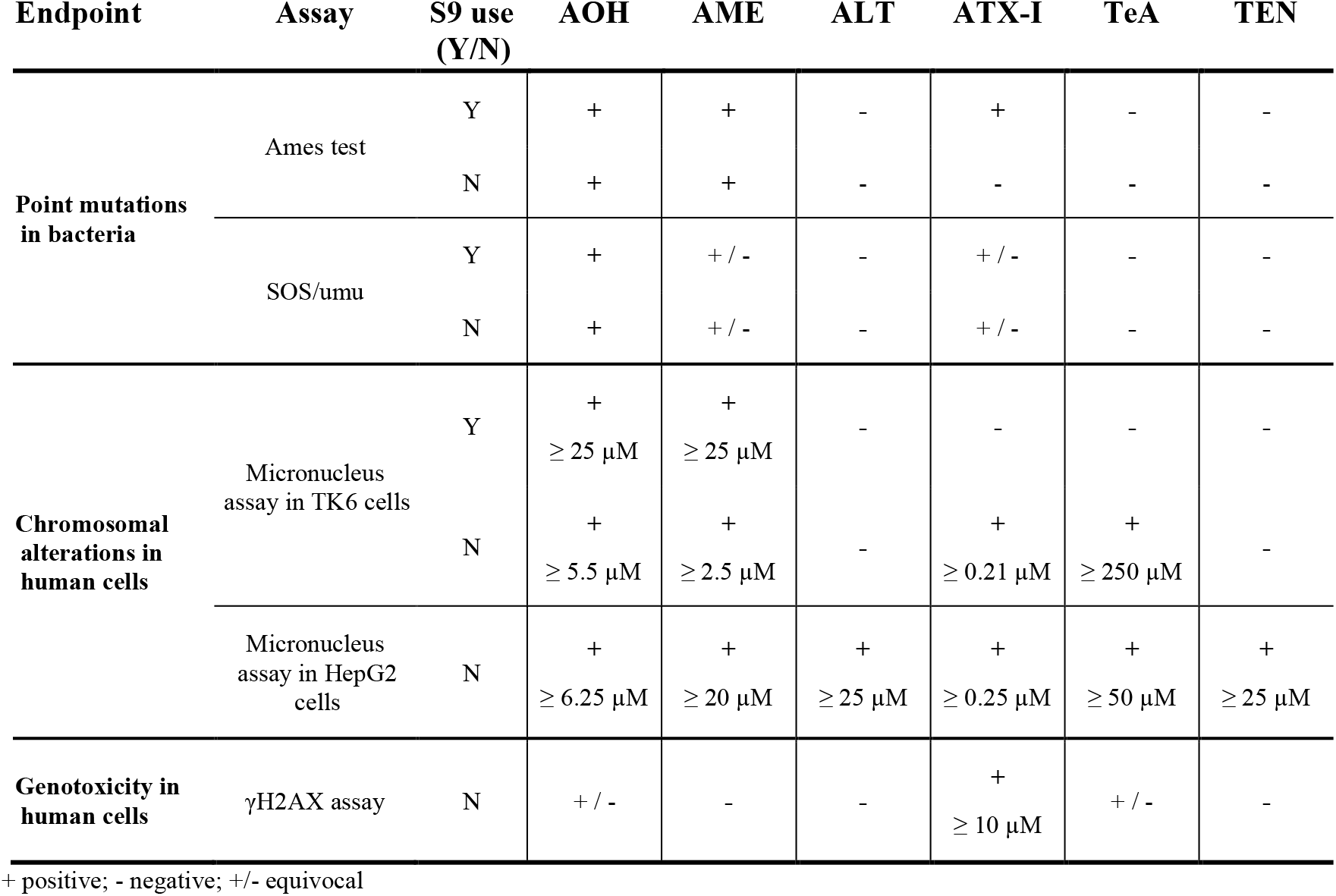
Overview of the outcomes of the genotoxicity assays.

## 4. Discussion

The aim of this work was to fill the data gaps using established toxicological tests, in line with OECD guidelines and the current recommended EFSA and WHO/FAO genotoxicity testing strategies (EFSA 2011a; WHO 2009). Under the coordinated project in PARC, the high-quality test samples from the same batch have been distributed to perform all testing, warranting comparability of data. The EFSA (2011a) Scientific Opinion recommends a two-tiered approach. The initial assessment, or Tier 1, consists of a core *in vitro* battery designed to cover three critical genetic endpoints: gene mutation, clastogenicity, and aneuploidy. This tier typically requires the Bacterial Reverse Mutation Test (Ames test; OECD TG 471) or the Mammalian Cell Gene Mutation (MCGM) assay (OECD TG 476), alongside the In Vitro Micronucleus Test (OECD TG 487). If the results of Tier 1 are negative, the substance is considered to have no genotoxic potential, and no further testing is required. However, if results are equivocal, additional testing may be necessary for clarification (EFSA 2011a; WHO 2009). A positive result in any Tier 1 assay triggers the requirement for Tier 2 (*in vivo*) testing to determine if the genotoxic activity is expressed in a whole-organism model. The choice of the *in vivo* follow-up is guided by the specific endpoint identified in Tier 1 and the requirement for adequate target tissue exposure. Recommended assays include the Mammalian Erythrocyte Micronucleus Test (OECD TG 474), the Transgenic Rodent Gene Mutation Assay (OECD TG 488), or the In Vivo Mammalian Alkaline Comet Assay (OECD TG 489). A negative result in Tier 2 generally concludes the assessment, indicating no genotoxic hazard. Conversely, positive results in any Tier 2 test identify a definitive genotoxic hazard, usually precluding the need for further testing (such as germ cell assays) regardless of carcinogenicity data.

The present publication covers *in vitro* testing aligned with the current EFSA testing strategy. In general, it is considered that two tests, normally Ames and MN, provide sufficient information, as they cover the three main genotoxicity endpoints (Ames test: mutagenicity and MN: clastogenicity and aneuploidy). To further support the current shift toward mechanistic data and reduced animal testing, the present study includes additional assays that facilitate the unraveling of the mode of action of these contaminants. Such data can contribute for the development of New Approach Methodologies (NAMs) integrated within Adverse Outcome Pathways (AOPs).

For instance, for the apical endpoint for mutagenicity, the SOS/umu test was also used. Although is not regulatory compliant, the test was selected because a high degree of agreement with the standardized Ames test (OECD TG 471) has been found (Reifferscheid and Heil 1996), and because it is an assay that requires a much smaller quantity of the test product than the Ames test. Validating this type of assay is important since the availability of a sufficient quantity of toxin is the bottleneck in mycotoxin evaluation, as these substances are either very expensive, or, in the case of emerging ones, are not commercially available in sufficient quantities. It is a medium-throughput assay for genotoxicity screening (Oda et al. 1985; Reifferscheid et al. 1991), which allows us to evaluate the DNA damaging effect of several compounds (six per 96-well plate). It needs relatively small amounts of test item (around 2 mg), and results are available within two days. The test is carried out in *S.Typhimurium* TA1535/pSK1002 with and without metabolic activation (S9 mix). *S.Typhimurium* TA1535 incorporates a pSK1002 plasmid containing the umuC gene fused to a lacZ reporter gene. The umuC gene is activated as part of the bacterial SOS response. In turn, this promotes the β-galactosidase activity associated with lacZ, which is assessed by a colorimetric reaction. This assay has been recently applied to regulated and emerging mycotoxins and has demonstrated a high degree of concordance with the known genotoxicity of these mycotoxins (Al-Ayoubi et al. 2023; Alonso-Jauregui et al. 2021; Alonso-Jauregui et al. 2022).

In order to detect chromosomal aberrations, the In Vitro Mammalian cell Micronucleus Test (OECD TG 487) was performed using two distinct cell lines, TK6 and HepG2, complemented by FISH analysis. While TK6 is one of the standard OECD-recommended cell lines for this assay, HepG2 cells were included to cover potential effects in liver, which has been recognized as target organ of some *Alternaria* toxins (Peach et al. 2024). In fact, the use of HepG2 cells is also a possibility depicted in the TG 487 and is justified based on their demonstrated performance in the test by the same authors (Dias et al. 2014; Pinto et al. 2014; Vasconcelos et al. 2019). This is also relevant in the context of the EFSA genotoxicity testing strategy, where the *in vivo* micronucleus assay in non-erythropoietic cells (e.g. liver and gastrointestinal tract) may be recommended at Tier 2, particularly when an assessment of *in vivo* aneugenicity is required. Furthermore, FISH analysis was integrated into the TK6 experiments to effectively differentiate between clastogenicity (structural damage) and aneugenicity (numerical chromosome loss or gain leading to aneuploidy).

Finally, the γH2AX assay was incorporated as a sensitive biomarker for the rapid detection of DNA double-strand breaks, providing mechanistic insight into the early cellular response to genotoxic stress. It is currently under consideration to be included as an OECD Test Guidelines (OECD 2026).

Among five tested bacterial strains (TA97a, TA98, TA100, TA102, and TA1535) AOH induced a > 2-fold increase in revertant colony number in strain TA97a (irrespective of metabolic activation) and in TA102 only in the presence of metabolic activation (at 0.03165 mg/plate). In contrast, Schrader et al. (2006) reported that AOH at nontoxic concentrations did not cause a > 2-fold increase in revertant colonies in strain TA97a (regardless of metabolic activation system) with observed toxicity at ≥ 0.05 and ≥ 0.1 mg/plate without and with S9, respectively. Similarly, Davis and Stack (1994) reported negative results for tested two strains TA98 and TA100 even at concentrations 7.5-fold higher (0.75 mg/plate) than those used in the present study, both with and without S9. There was a weak increase (IF < 2) in TA100 (with and without S9) and TA104 (with S9) induction factors tested up to 0.1 mg/plate without observed toxicity (Schrader et al. 2006; Schrader et al. 2001). Furthermore, Schrader et al. (2006) demonstrated that AOH at nontoxic concentrations induced a dose-dependent increase in average number of revertants in TA102 without (0.05 mg/plate) and with S9 (≥ 0.05 mg/plate), reaching a 2- to 3-fold increase, with observed toxicity at 0.1 mg/plate in the absence of S9. In our study, toxicity was observed at 0.1 mg/plate regardless of S9, which may explain the absence of a clear mutagenic response without metabolic activation. Complementing the Ames data, AOH produced a clearly positive result in the SOS/umu assay, both in the presence and absence of metabolic activation, supporting its mutagenic potential in bacterial systems.

In mammalian cells, clastogenic activity of AOH becomes more apparent. In the *in vitro* MN assay using TK6 cells, AOH exhibited a concentration-dependent increase in MN formation both with and without S9. FISH analysis after 24h exposure without metabolic activation, suggested that AOH acts as a clastogen by causing structural chromosome aberrations, which is supported by the study of Solhaug et al. (2016). The MN results align with previous *in vitro* MN studies, in which AOH also induced chromosomal damage in three other cell types (i.e., RAW264.7, Ishikawa, and V79) (Lehmann et al. 2006; Solhaug et al. 2013). Nevertheless, in these previous studies, the assays were only performed without metabolic activation. In contrast, an *in vivo* MN assay in male Sprague-Dawley rats orally exposed to AOH (5.51, 11.03, and 22.05 µg/kg bw) for 28 consecutive days did not reveal chromosome damage in peripheral blood and liver (Miao et al. 2022). A recent review of Louro et al. (2024) highlighted the hypothesis suggesting that the absence of a genotoxic effect *in vivo* could be attributed to the low systemic bioavailability of AOH (Miao et al. 2022) or to rapid biotransformation and glutathione conjugation, as suggested by Pfeiffer et al. (2009) and Tiessen et al. (2013).

The capacity of AOH to induce MN in mammalian cells was further evaluated in HepG2 cells. AOH showed a sharp decline in HepG2 cell viability above 25 μM and an IC_50_ of 116.8 μM, consistent with previous hepatic model data (Mahmoud et al. 2022). Genotoxic effects in HepG2 cells were pronounced, with a 4.5-fold increase in the frequency of MNBNC over control and showing cytostatic activity at the highest concentration tested (50 µM), as assessed by a decrease of the CBPI. AOH consistently increased both NBUDs and NPBs frequencies, as well. Mechanistically, AOH has been shown to form reactive oxygen species and to interact with DNA topoisomerase, thereby generating both single- and double-strand DNA beaks (Solhaug et al. 2016). While, to our best knowledge, CBMN data on AOH in HepG2 cells are novel, our findings agree with previous studies using the alkaline unwinding assay and the γH2AX assay in hepatic and other mammalian cells, showing that AOH induces DNA strand breaks at comparable concentrations (≥ 12.5 μM) (Fleck et al. 2014; Hessel-Pras et al. 2019; Pfeiffer et al. 2007a).

Consistent with these findings, AOH also induced γH2AX phosphorylation in HepaRG cells, albeit at higher concentrations (300 µM) than those reported for HepG2 cells (Hessel-Pras et al. 2019). This shift in effective concentration may reflect the higher metabolic capacity of HepaRG cells compared to HepG2 cells, resulting in enhanced biotransformation and partial detoxification of AOH, thereby higher concentrations of AOH are necessary to elicit detectable DNA damage.

AME induced a dose-dependent increase in revertant counts, exceeding a 2-fold induction in strain TA97a (irrespective of metabolic activation) at concentrations as low as 0.0086 mg/plate. In contrast, Schrader et al. (2006) reported no increase in revertant colonies in TA97a even at concentrations up to 0.1 mg/plate. However, they observed a weak mutagenic response (IF < 2) in strains TA102 (with and without S9) and TA104 (with S9). In our study, 0.1 mg/plate was already toxic to TA102 in the absence of S9, potentially limiting observable mutagenicity under these conditions. A strong mutagenic response has previously been reported in *E. coli* strain ND-160 without S9, with IF of approximately 5 and 10 at 0.05 and 0.1 mg/plate, respectively (An et al. 1989). Consistent with our findings, the absence of mutagenic activity in strains TA98 and TA100 has been confirmed by other studies even at concentrations up to 7.5-fold higher than those tested in our work (Davis and Stack 1994; Schrader et al. 2001). In addition, AME was also negative in strains TA1537 and TA1538 (Davis and Stack 1994). Contradictory results for AME could be due to the presence of trace amounts of highly mutagenic altertoxins in their toxin samples, which were purified from cultures, whereas in the present study chemically synthesized AME was used. AME also yielded a weak positive response in the SOS/umu assay (IF > 1.5 but without a clear dose-response, with and without S9), supporting the DNA-damaging potential of AME in bacteria but suggesting limited potency compared to other *Alternaria* toxins.

In mammalian systems, genotoxic effects of AME were more pronounced. In the MN assay, AME induced a significant concentration-dependent increase in the frequency of MNBNCs in TK6 cells, both with and without S9 metabolic activation, after 3h and 24h exposure, respectively. The effect was more pronounced without metabolic activation, as an increase was observed already at 2.5 µM, whereas with metabolic activation the effect was observed from 25 µM onwards. Centromere staining and FISH analysis indicated a clastogenic mode of action. Among the previously published *in vitro* assays performed with AME, surprisingly no MN data were reported (Louro et al. 2024). In contrast, an *in vivo* MN assay is available in which male Sprague-Dawley rats received oral doses of AME (1.84, 3.67, or 7.35 µg/kg bw/day) for 28 days. In this study, an increase in MN formation was observed at the highest dose in both the bone marrow and peripheral blood, indicating that AME is capable of inducing chromosomal damage *in vivo* under sustained exposure (Tang et al. 2022).

In HepG2 cells, AME showed a dose-dependent cytotoxic effect above 25 μM (IC_50_ ≈ 20 μM), consistent with previous findings (den Hollander et al. 2022). AME induced a significant dose-dependent increase in MN frequency in HepG2 cells at concentrations above 20 μM. Previous work showed that AME induced DNA strand breaks in HepG2 cells but not in HT-29 cells, likely due to the HepG2 cells’ inability to metabolize AME into its conjugated form (Pfeiffer et al. 2007a). In fact, analysis of the exposure culture medium of HepG2 cells revealed that approximately 75% of AME remained in its unconjugated form, while AME was completely conjugated in HT-29 cells. These findings suggest that the strand-breaking activity is primarily caused by the parent toxin and that glucuronidation represents a metabolic inactivation pathway for the clastogenicity of this mycotoxin (Pfeiffer et al. 2007b).

This interpretation is further supported by our γH2AX findings. AME did not induce γH2AX phosphorylation in HepaRG cells up to 100 µM, suggesting efficient detoxification in this cell line. HepaRG cells possess functional DNA repair mechanisms and express a phase I and phase II metabolic enzyme profile which is close to that of primary human hepatocytes. Additionally, Hessel-Pras et al. (2019) showed that AME toxicity was reduced by pre-treatment with S9 liver homogenate, as indicated by decreased cytotoxicity and H2AX phosphorylation to control levels. The same study reported low cytotoxicity in HepaRG cells with or without additional metabolic activation, likely due to their active xenobiotic-metabolizing enzymes, which confer resilience (Hessel-Pras et al. 2019). Together, these findings suggest that the differentiation status of cells influences their sensitivity to *Alternaria* toxins and that metabolic detoxification may mitigate their effects.

ALT did not induce mutagenic response in *S.*Typhimurium strains TA97a, TA98, TA100, TA102, and TA1535 either in the absence or presence of a metabolic activation system, at concentrations up to 0.292 mg/plate. Similarly, negative Ames assays were reported in strains TA97, TA98, TA100, TA102, and TA104 up to 0.1 mg/plate, both with and without a metabolic activation system, using plate incorporation assay (Schrader et al. 2006; Schrader et al. 2001). ALT was likewise negative in the SOS/umu assay, suggesting absence of mutagenicity in bacterial systems.

Consistent with these findings, ALT did not induce a significant dose-dependent increase in MN formation in TK6 cells under any of the tested conditions (i.e. 3h with and without S9 and 24h without S9) in the *in vitro* MN assay. To date, no additional *in vitro* or *in vivo* MN assay has been conducted for ALT (Louro et al. 2024).

In contrast, ALT demonstrated clear genotoxic activity in HepG2 cells. Although cytotoxicity was low (IC_50_ = 286 μM), ALT induced a strong concentration-dependent increase in the frequency of MNBNCs and NBUDs. Previous comet assay data in human colon cells did not detect DNA damage, but this discrepancy likely reflects endpoint differences, as comet assays primarily detect single- and double-strand DNA breaks instead of chromosomal instability (Fehr et al. 2009). In addition, DNA damage detected by the comet assay often reflects reversible DNA lesions that are still amenable to repair, while the CBMN assay detects clastogenic and aneugenic effects (Fehr et al. 2009; Xiao et al. 2014). The apparent divergent results of ALT in HepG2 cells and TK6 cells can largely be attributed to fundamental differences in cell cycle control, genomic surveillance, and metabolic capacity between both cell types. TK6 cells possess an intact spindle assembly checkpoint and functional p53 signaling, which enables efficient detection of mitotic and DNA damage, promoting cell cycle arrest or mitochondrial-mediated apoptosis via Cdk1–Bax/Bak signaling (Darweesh et al. 2021; Honma 2005; Ruan et al. 2019; Whitwell et al. 2015). In contrast, HepG2 cells display dysregulation of multiple mitotic regulators, resulting in weakened checkpoint enforcement and allowing cells to progress through mitosis despite spindle defects (Carloni et al. 2018; Yang et al. 2022). This tolerance of mitotic errors, combined with compromised p53 function (Bressac et al. 1990; Li et al. 2024), promotes the persistence of genetically damaged cells. Consequently, TK6 cells tend to eliminate heavily damaged cells, whereas HepG2 cells may continue proliferating, thereby increasing the likelihood of micronucleus formation and other nuclear abnormalities. Notably, ALT induced the highest frequency of atypical nuclear morphologies including FUS, CIR, and HS nuclei supporting that this mycotoxin may destabilize nuclear architecture and interfere with spindle function. These nuclear anomalies, previously associated with chromosomal instability under folate deficiency, appear to originate from persistent nuclear strands and incomplete chromatid separation during mitosis (Bull et al. 2012; Fenech 2020; Lee et al. 2014). Supporting this hypothesis, previous studies have shown that some *Alternaria* mycotoxins can disrupt mitotic spindle assembly and cytokinesis, leading to abnormal spindles, nuclear bridges, and irregular nuclear morphologies (Lehmann et al. 2006; Solhaug et al. 2013). Taken together, these observations suggest that spindle disruption and impaired chromosome segregation may contribute to the abnormal nuclear morphologies observed. Furthermore, longer exposure periods commonly applied in HepG2 assays increase the probability that indirect or pre-mitotic damage persists until the first nuclear division, thereby favoring micronucleus expression (Khoury et al. 2016; Sobol et al. 2012; Speit et al. 2012).

Interestingly, ALT was negative in the γH2AX assay, suggesting that its genotoxicity may not primarily involve double-strand break induction but rather mitotic spindle disruption and chromosomal instability. These cell differences underscore the importance of using multiple *in vitro* models when assessing genotoxicity.

The perylene quinone ATX-I showed a strain-specific mutagenicity in the Ames test, inducing a > 2-fold increase at 0.01 mg/plate only in strains TA97a and TA102 without S9, suggesting that ATX-I may be metabolized into derivatives that are not mutagenic at this concentration. Schrader et al. (2006) similarly reported a mutagenic response in TA102 at 2 times lower concentration (0.005 mg/plate) without S9 and additionally at 5 times higher concentration than used in our study (0.05 mg/plate) in the presence of S9. In contrast, the same study classified ATX-I as a weak mutagen for TA97a, regardless of metabolic activation, even at concentrations 5 times higher than those used in our study. ATX-I was also identified as a weak mutagen for TA104 (Schrader et al. 2006) and as mutagenic for TA1537 (Stack and Prival 1986). Literature data for TA98 and TA100 are inconsistent: Stack and Prival (1986) reported strong mutagenicity of ATX-I (up to 0.06 mg/plate) for both strains regardless of S9, whereas Schrader et al. (2001) later observed mutagenicity only in TA98 with S9 and not in TA100 under any condition, despite testing higher concentrations (up to 0.1 mg/plate). In the SOS/umu assay, ATX-I also tested positive, both without and with metabolic activation, although the response was weaker (IF > 1.5 but without a clear dose-response) and was classified as weak positive.

ATX-I significantly increased the MN formation in TK6 cells, but only in the absence of the S9 fraction. Interestingly, this increase became significant at 0.21 µM and appeared to reach a maximum in terms of MN formation from 0.41 µM, beyond which it did not increase further with increasing concentrations. The FISH analysis showed a majority of micronuclei without centromere for the four highest concentrations tested (from 0.41 µM to 8.3µM) suggesting a clastogenic mode of action rather than aneugenic. To date, only a few studies have been conducted to assess the genotoxicity of ATX-I and, specifically, no *in vitro* MN assay was performed previously for ATX-I. Furthermore, in the only available *in vivo* study, no genotoxic effect was observed using the micronucleus assay on peripheral blood and bone marrow of male Sprague-Dawley rats exposed for 28 days at up to 5.51 µg/kg bw/day (Zhu et al. 2022).

In HepG2 cells, ATX-I is the most hazardous mycotoxin tested, showing cytotoxicity even at low concentrations (IC_50_ = 56.8 μM), which suggests higher cytotoxicity than previously reported (Mahmoud et al. 2022). Genotoxic effects of ATX-I were evident in HepG2 cells, even though solubility limitations restricted testing above 1 μM. The significant induction of MN and NBUDs observed in the present work is consistent with previous reports using the alkaline unwinding assay in V79, HepG2, and Caco-2 cells that detected DNA strand breaks (Fleck et al. 2014). ATX-I was also able to induce the phosphorylation status of γH2AX in HepaRG cells, demonstrating its ability to induce primary DNA damage.

TeA did not induce mutagenic response in *S.*Typhimurium strains TA98 and TA100 without and with metabolic activation system up to 2.5 mg/plate. Similarly, negative Ames assays were reported in strains TA97, TA98, TA100, TA102 and TA104 up to 0.1 mg/plate with and without metabolic activation system, using plate incorporation assay (Schrader et al. 2006; Schrader et al. 2001). TeA was also negative in the SOS/umu assay, supporting the conclusion that TeA is not mutagenic in bacterial systems.

In mammalian cells, evidence for genotoxicity was limited and mainly observed at high concentrations and prolonged exposure. In TK6 cells, TeA induced DNA damage only after 24h exposure without metabolic activation and at relatively high concentrations (≥ 250 µM). The proportion of MN CENT+ and MN CENT-obtained after the FISH staining, were similar to the clastogenic positive control (i.e, MMS) and indicated the ability to induce structural chromosome aberration (clastogenicity) rather than numerical chromosome aberration (aneugenicity). Notably, this represents the first MN dataset for TeA, as neither *in vitro* nor *in vivo* MN data were available to date for TeA (Louro et al. 2024).

In HepG2 cells, TeA exhibited moderate cytotoxicity with an IC_50_ of 446.31 μM, consistent with previous reports, although values vary depending on exposure duration (den Hollander et al. 2022; Mahmoud et al. 2022). After 48h exposure, TeA revealed a genotoxic effect in HepG2 cells, with a significant increase in the frequency of micronucleated BNC at concentrations equal to or higher than 50 μM. Toxicokinetic findings from *in vivo* studies suggest, that TeA undergoes limited metabolism, as the parent mycotoxin is largely excreted unchanged in the urine of rats and mice and no urinary metabolites have been identified (Asam et al. 2013; Puntscher et al. 2019a). Mechanistically, several studies have suggested that a potential mechanism for TeA-induced MN formation may involve the generation of ROS, leading to DNA strand breakage and subsequently to chromosomal breakage (Higashi 1988; Sandermann Jr 1988; Zhou and Qiang 2008).

Consistently with this interpretation, γH2AX induction in HepaRG cells was observed only at extremely high concentrations of 1000 µM. Such concentrations are far above those expected under realistic exposure scenarios, raising questions about the biological relevance of this result and suggesting that indirect mechanisms may rather play a role.

TEN did not induce mutagenicity in the Ames test in any of the tested *S.*Typhimurium strains (TA97a, TA98, TA100, TA102 and TA1535) up to 0.415 mg/plate, irrespective of metabolic activation. Similarly, Schrader et al. (2001, 2006) reported negative Ames assay in strains TA97, TA98, TA100, TA102 and TA104 up to 0.1 mg/plate with and without metabolic activation system, using plate incorporation assay. TEN was also negative in the SOS/umu assay, further supporting the absence of mutagenicity in bacterial systems.

In the MN assay using TK6 cells, TEN did not induce a concentration-dependent increase in chromosome damage under any of the tested conditions. To our knowledge, TEN was not previously tested with the MN assay, and no *in vivo* study on TEN is available so far. However, a comet assay revealed negative findings in HEK293T cells exposed for 24h to 25 µM without metabolic activation supporting a lack of primary DNA damage in short-term assays (Tran et al. 2020).

In contrast, prolonged exposure (48h) to TEN revealed weak genotoxic effects in hepatic cells. In HepG2 cells, TEN displayed very low cytotoxicity with only slight reductions in viability at concentrations above 200 μM and an extrapolated IC_50_ far above levels expected in human blood following dietary exposure. These findings are in line with those from previous cytotoxicity studies (Hessel-Pras et al. 2019; Tran et al. 2020). Despite this low cytotoxic profile, a dose-dependent increase in MN frequency was observed after 48h exposure. The positive genotoxicity in HepG2 cells, and not in TK6 cells, can again be explained by the differences in checkpoint regulation and p53 signaling of both cell lines, as well as the exposure duration, already discussed in the ALT section.

Consistent with the generally weak genotoxic profile, TEN was negative in the γH2AX in HepaRG cells after 24h exposure, similar to findings in a γH2AX assay in HepG2 cells exposed for 4h (Hessel-Pras et al. 2019).

### Summary of the results

The present study provides a comprehensive evaluation of the genotoxic potential of selected *Alternaria* toxins (AOH, AME, ALT, ATX-I, TeA, and TEN) across bacterial and mammalian test systems. Among the tested toxins, AOH and ATX-I exhibited clear effects regarding genotoxicity and mutagenicity. Both compounds induced mutagenic responses in bacterial systems and demonstrated pronounced clastogenicity in mammalian cells by increasing MN formation in TK6 and HepG2 cells and inducing γH2AX phosphorylation. For ATX-I, genotoxic effects were observed at particularly low micromolar concentrations, showing its high potency compared to the other tested *Alternaria* toxins. AME showed clear mutagenicity in strain TA97a and induced MN formation in TK6 and HepG2 cells, with a clastogenic mode of action. In contrast to AOH and ATX-I, it seems that the genotoxic effect of AME is highly dependent on biotransformation, as detoxification processes mitigate its effect. ALT and TEN were negative in bacterial systems and did not induce MN formation in TK6 cells. However, both toxins produced genotoxic effects in HepG2 cells after prolonged exposure, suggesting that spindle interferences and chromosomal instability, rather than direct induction of DNA double-strand breaks may play a role specifically in liver cells. This finding is consistent with the absence of γH2AX induction. Overall, TeA showed the least evident genotoxic activity, since it was negative in bacterial assays and induced MN formation only at high concentrations and extended exposure in mammalian cells. γH2AX induction occurred only at extremely high concentrations, raising questions regarding the biological relevance in realistic exposure scenarios.

Since all tested *Alternaria* toxins have been shown to be genotoxic *in vitro* in at least one endpoint, the next step in most regulatory contexts would be to confirm *in vivo* whether the observed adverse effects also occur in the whole organism. Before embarking on animal studies, additional relevant information on the substance should be integrated, including chemical reactivity that may predispose to site-of-contact effects, bioavailability, metabolism, toxicokinetics, and potential target organ specificity. This information enhancing the experimental design, ensuring that new *in vivo* studies address the genotoxic endpoint identified as positive *in vitro* and focus on appropriate target organs or tissues. It is recognized that genotoxic chemicals cannot be deliberately added to food at any dose level. If present as unavoidable contaminants, risk assessment (RA) approaches based on the Margin of Exposure (MOE) may be used to advise risk managers on the level of concern (Bennekou et al. 2025). This MOE represents the ratio between a reference point derived from animal studies and the estimated human intake. Since genotoxic carcinogens are generally assumed to have no safe threshold, meaning that any exposure may carry some risk, the MOE approach or alternative approaches based on the principle As Low As Reasonably Achievable (ALARA) should be considered. Likewise, when a food contaminant is identified as both genotoxic and carcinogenic, the Joint FAO/WHO Expert Committee on Food Additives (JECFA) generally concludes that no recognizable threshold exists below which the substance would be considered safe (Barlow et al. 2006). Consequently, JECFA does not establish a traditional acceptable daily intake or provisional tolerable weekly intake, as any level of exposure may potentially pose a risk.

## Statements and Declarations

The authors declare that they have no conflict of interest. During the preparation of this work, the authors used ChatGPT for grammar and language editing. After using this tool, the authors reviewed and edited the content as needed and take full responsibility for the content of the publication.

## Acknowledgments

The European Partnership for the Assessment of Risks from Chemicals (PARC) has received funding from the European Union’s Horizon Europe research and innovation program under Grant Agreement No 101057014 and has received co-funding of the authors’ institutions. Views and opinions expressed are, however, those of the author(s) only and do not necessarily reflect those of the European Union or the Health and Digital Executive Agency. Neither the European Union nor the granting authority can be held responsible for them.

AV thanks the Grant PID2021-126026OB-I00-MYCOCANCER funded by the Spanish Ministry of Science and Innovation MICIU/AEI/ 10.13039/501100011033 and by ERDF/EU.

Sciensano’s contribution was co-funded by the Belgian Federal Public Service (FPS) through the grant received for the project RF 23/18 ENNIATOX.

