## Supplemental material for "Hazard characterization of *Alternaria* toxins – filling data gaps on *in vitro* genotoxicity"

### Bacterial Reverse Mutation Assay

Table S 1: Mutagenic activity of *Alternaria* toxins in *S.Typhimurium* in the absence (−S9) or presence (+S9) of an external metabolizing enzyme system (10% rat liver S9 mix).

| Treatment (mg/plate) | Salmonella typhimurium strain | | | | | | | | | |
|---|---|---|---|---|---|---|---|---|---|---|
| | TA97a | | TA98 | | TA100 | | TA102 | | TA1535 | |
| | -S9 | +S9 | -S9 | +S9 | -S9 | +S9 | -S9 | +S9 | -S9 | +S9 |
| *AOH* | | | | | | | | | | |
| NC | 192.0 ± 15.7 | 217.0 ± 3.6 | 40.0 ± 2.0 | 33.3 ± 4.2 | 93.0 ± 3.6 | 119.0 ± 3.6 | 431.0 ± 14.2 | 418.7 ± 25.7 | 10.7 ± 1.2 | 7.0 ± 1.7 |
| SC | 189.3 ± 15.7 | 238.7 ± 9.5 | 34.3 ± 3.1 | 29.0 ± 7.5 | 89.0 ± 5.6 | 103.3 ± 13.9 | 393.3 ± 6.7 | 373.7 ± 14.2 | 9.0 ± 1.0 | 8.7 ± 1.5 |
| 0.001 | 332.7 ± 28.5 | 232.7 ± 15.8 | 37.0 ± 3.0 | 26.7 ± 3.1 | 76.0 ± 7.5 | 93.0 ± 4.4 | 339.7 ± 21.6 | 383.0 ± 25.5 | 6.0 ± 1.7 | 10.7 ± 2.3 |
| 0.00317 | **428.0 ± 21.0** | 309.3 ± 19.0 | 34.0 ± 5.6 | 26.3 ± 2.1 | 86.0 ± 7.2 | 98.7 ± 4.0 | 406.0 ± 16.7 | 404.7 ± 17.5 | 7.7 ± 1.5 | 11.7 ± 3.2 |
| 0.01001 | **652.0 ± 39.1** | 457.7 ± 22.7 | 39.7 ± 8.0 | 32.7 ± 4.9 | 83.0 ± 3.6 | 100.3 ± 4.0 | 372.3 ± 42.3 | 425.0 ± 18.7 | 15.0 ± 4.4 | 13.3 ± 3.8 |
| 0.03165 | **877.7 ± 40.8** | 741.7 ± 16.5 | 47.3 ± 2.3 | 30.3 ± 2.1 | 109.7 ± 5.5 | 131.7 ± 8.5 | 634.0 ± 21.5 | **753.3 ± 33.7** | 13.0 ± 1.7 | 9.7 ± 1.5 |
| 0.1 | 97.0 ± 22.9[b] | 667.7 ± 33.9 | 50.3 ± 10.0 | 16.7 ± 2.5[a] | 102.3 ± 6.5[a] | 136.0 ± 4.6[a] | 216.7 ± 18.0[b] | 1067.0 ± 86.1[b] | 5.3 ± 1.2[b] | 1.3 ± 0.6[b] |
| PC | **689.3 ± 45.1** | **846.0 ± 30.8** | **300.0 ± 13.5** | **298.3 ± 14.7** | **283.0 ± 10.5** | **641.7 ± 33.5** | **972.0 ± 23.6** | **1061.3 ± 124.5** | **193.0 ± 15.0** | **147.3 ± 22.0** |
| *AME* | | | | | | | | | | |
| NC | 237.3 ± 35.1 | 239.3 ± 24.6 | 23.0 ± 1.0 | 33.3 ± 5.0 | 82.3 ± 4.0 | 97.0 ± 9.5 | 408.7 ± 16.3 | 499.0 ± 47.8 | 10.3 ± 2.5 | 12.3 ± 4.6 |
| SC | 236.0 ± 43.7 | 210.7 ± 37.3 | 20.7 ± 1.5 | 32.0 ± 10.4 | 72.0 ± 11.5 | 88.0 ± 13.9 | 388.3 ± 28.7 | 430.3 ± 32.3 | 11.7 ± 3.2 | 10.7 ± 2.1 |
| 0.0027 | 333.7 ± 33.5 | 306.7 ± 59.2 | 24.7 ± 1.2 | 33.3 ± 9.7 | 74.3 ± 11.4 | 92.7 ± 14.2 | 406.0 ± 24.9 | 499.7 ± 36.1 | 9.7 ± 2.5 | 9.0 ± 5.6 |
| 0.0086 | **497.0 ± 32.4** | **453.3 ± 95.0** | 17.3 ± 5.5 | 37.3 ± 11.2 | 77.0 ± 8.9 | 104.7 ± 13.2 | 410.7 ± 23.1 | 511.3 ± 35.6 | 10.0 ± 1.7 | 8.0 ± 3.6 |
| 0.027 | **723.0 ± 111.2** | **769.3 ± 141.1** | 17.7 ± 9.1 | 42.3 ± 6.5 | 87.3 ± 10.2 | 110.0 ± 13.1 | 431.0 ± 17.0 | 501.7 ± 31.5 | 11.0 ± 2.6 | 8.0 ± 1.0 |
| 0.086 | **798.0 ± 121.7** | **872.3 ± 196.3** | 26.3 ± 5.7 | 39.3 ± 7.8 | 77.7 ± 2.3 | 109.0 ± 23.8 | 375.7 ± 45.1 | 544.7 ± 123.3 | 10.3 ± 2.1 | 10.7 ± 3.2 |
| 0.272 | **844.3 ± 86.3** | **975.3 ± 288.8** | 13.7 ± 4.5 | 40.3 ± 5.5 | 78.3 ± 11.6 | 107.7 ± 17.2 | 165.0 ± 34.2 | 266.0 ± 66.2 | 8.3 ± 2.9 | 6.7 ± 1.5 |
| PC | **602.7 ± 86.2** | **630.0 ± 22.3** | **341.3 ± 67.9** | **133.7 ± 15.6** | **351.7 ± 35.0** | **505.7 ± 35.0** | **1441.33 ± 102.81** | **1439.3 ± 118.1** | **121.0 ± 54.1** | **95.7 ± 7.2** |
| *ALT* | | | | | | | | | | |
| NC | 188.0 ± 17.1 | 228.3 ± 4.5 | 32.0 ± 4.0 | 36.7 ± 5.1 | 88.7 ± 16.8 | 104.7 ± 17.0 | 384.0 ± 8.2 | 467.0 ± 22.7 | 10.0 ± 3.0 | 11.7 ± 4.9 |
| SC | 176.0 ± 13.0 | 227.7 ± 25.7 | 30.3 ± 3.2 | 31.7 ± 6.4 | 95.7 ± 13.3 | 100.7 ± 16.3 | 388.3 ± 28.7 | 430.3 ± 32.3 | 13.7 ± 2.9 | 12.0 ± 2.0 |
| 0.0029 | 182.3 ± 4.7 | 231.7 ± 20.5 | 24.0 ± 2.0 | 31.7 ± 6.0 | 78.0 ± 11.5 | 96.0 ± 12.5 | 392.3 ± 21.1 | 436.0 ± 18.0 | 11.3 ± 3.1 | 15.0 ± 2.6 |
| 0.0092 | 178.7 ± 13.3 | 263.3 ± 20.6 | 26.3 ± 5.5 | 29.3 ± 2.1 | 81.3 ± 15.9 | 87.3 ± 21.0 | 385.0 ± 24.2 | 458.7 ± 63.1 | 13.3 ± 3.5 | 9.0 ± 2.0 |
| 0.029 | 199.7 ± 13.8 | 246.3 ± 11.0 | 32.0 ± 6.1 | 31.0 ± 3.0 | 83.3 ± 18.4 | 86.0 ± 7.00 | 354.7 ± 33.0 | 457.7 ± 50.1 | 9.7 ± 2.1 | 10.0 ± 4.0 |
| 0.092 | 186.0 ± 13.0 | 237.0 ± 37.3 | 29.7 ± 6.1 | 25.0 ± 4.4 | 101.0 ± 21.6 | 85.3 ± 11.1 | 317.3 ± 35.9 | 361.7 ± 72.8 | 10.7 ± 1.5 | 14.7 ± 4.0 |
| 0.292 | 169.0 ± 21.0 | 243.7 ± 34.2 | 34.0 ± 8.7 | 36.0 ± 5.2 | 72.7 ± 2.5 | 98.3 ± 23.5 | 191.3 ± 56.2 | 281.3 ± 44.0 | 11.3 ± 2.1 | 12.7 ± 4.0 |
| PC | **544.7 ± 27.2** | **612.3 ± 48.2** | **307.3 ± 11.1** | **121.7 ± 16.3** | **430.7 ± 192.6** | **880.0 ± 153.0** | **1468.0 ± 177.2** | **1622.7 ± 184.3** | **143.7 ± 11.6** | **102.3 ± 9.6** |
| *ATX1* | | | | | | | | | | |
| NC | 211.7 ± 11.4 | 251.7 ± 10.2 | 19.7 ± 2.5 | 36.3 ± 5.7 | 90.0 ± 7.0 | 114.3 ± 12.1 | 360.7 ± 10.5 | 397.3 ± 15.6 | 7.3 ± 0.6 | 9.3 ± 1.5 |
| SC | 222.3 ± 5.9 | 315.7 ± 8.4 | 20.0 ± 1.7 | 32.0 ± 4.6 | 73.3 ± 2.1 | 104.0 ± 7.2 | 363.0 ± 9.2 | 424.3 ± 16.2 | 9.3 ± 1.2 | 10.0 ± 1.0 |

| | | | | | | | | | | |
|---|---|---|---|---|---|---|---|---|---|---|
| 0.0001 | 268.7 ± 17.2 | 318.7 ± 7.1 | 23.0 ± 3.0 | 28.0 ± 2.6 | 90.0 ± 7.0 | 87.7 ± 5.9 | 380.0 ± 13.5 | 449.7 ± 14.1 | 9.3 ± 1.5 | 9.0 ± 1.0 |
| 0.0003169 | 255.7 ± 10.0 | 292.0 ± 17.6 | 21.7 ± 3.5 | 29.3 ± 4.0 | 80.7 ± 4.0 | 103.7 ± 9.3 | 379.3 ± 16.3 | 420.3 ± 12.5 | 11.3 ± 1.5 | 9.0 ± 1.0 |
| 0.001 | 273.7 ± 12.1 | 311.3 ± 22.1 | 21.0 ± 4.4 | 37.0 ± 3.6 | 75.3 ± 4.5 | 98.7 ± 9.0 | 364.3 ± 16.3 | 424.0 ± 7.8 | 8.0 ± 1.0 | 13.0 ± 2.0 |
| 0.00316 | 318.0 ± 15.1 | 324.3 ± 9.3 | 31.7 ± 4.2 | 33.0 ± 1.7 | 93.0 ± 12.5 | 102.0 ± 7.8 | 483.3 ± 18.6 | 419.3 ± 19.5 | 7.3 ± 0.6 | 8.3 ± 1.5 |
| 0.010 | **451.7 ± 10.5** | 352.3 ± 11.0 | 31.7 ± 3.2 | 33.7 ± 5.1 | 100.3 ± 7.0 | 108.7 ± 10.3 | **809.3 ± 27.0** | 473.7 ± 20.1 | 12.3 ± 2.1 | 12.3 ± 0.6 |
| PC | **683.0 ± 34.6** | **892.7 ± 34.7** | **303.0 ± 15.5** | **180.3 ± 9.0** | **344.0 ± 12.5** | **676.0 ± 38.7** | **1014.0 ± 34.6** | **1360.3 ± 51.9** | **131.0 ± 17.3** | **227.3 ± 8.7** |
| *TeA* | | | | | | | | | | |
| NC | NA | NA | 19.7 ± 2.1 | 44.3 ± 7.0 | 83.7 ± 5.5 | 103.0 ± 7.6 | NA | NA | NA | NA |
| SC | NA | NA | 20.7 ± 1.5 | 42.7 ± 5.0 | 89.7 ± 4.2 | 98.3 ± 8.7 | NA | NA | NA | NA |
| 0.025 | NA | NA | 22.3 ± 1.5 | 50.0 ± 4.4 | 96.7 ± 7.1 | 94.7 ± 7.2 | NA | NA | NA | NA |
| 0.079 | NA | NA | 23.0 ± 2.6 | 46.0 ± 4.6 | 87.7 ± 9.6 | 119.3 ± 5.7 | NA | NA | NA | NA |
| 0.25 | NA | NA | 20.3 ± 1.2 | 51.7 ± 3.1 | 84.0 ± 10.0 | 114.0 ± 13.1 | NA | NA | NA | NA |
| 0.791 | NA | NA | 20.3 ± 2.1 | 45.7 ± 2.1 | 100.0 ± 13.5 | 90.3 ± 8.0 | NA | NA | NA | NA |
| 2.5 | NA | NA | 22.7 ± 3.2 | 41.3 ± 3.2 | 98.3 ± 14.7 | 106.3 ± 8.6 | NA | NA | NA | NA |
| PC | NA | NA | **163.7 ± 12.1** | **203.0 ± 19.0** | **296.3 ± 31.6** | **757.7 ± 36.5** | NA | NA | NA | NA |
| *TEN* | | | | | | | | | | |
| NC | 189.7 ± 19.8 | 228.3 ± 4.5 | 34.0 ± 5.3 | 36.7 ± 5.1 | 88.7 ± 16.8 | 104.7 ± 17.0 | 426.7 ± 13.9 | 467.0 ± 22.7 | 10.0 ± 3.0 | 12.3 ± 5.5 |
| SC | 176.0 ± 13.0 | 241.0 ± 15.7 | 32.3 ± 3.8 | 31.7 ± 6.4 | 104.7 ± 22.6 | 104.7 ± 12.7 | 404.3 ± 23.9 | 430.3 ± 32.3 | 13.7 ± 2.9 | 12.7 ± 2.3 |
| 0.0041 | 173.0 ± 9.2 | 234.0 ± 16.7 | 28.0 ± 7.5 | 34.3 ± 10.1 | 99.7 ± 11.2 | 90.3 ± 18.7 | 371.0 ± 26.9 | 420.3 ± 17.2 | 11.0 ± 2.0 | 14.7 ± 2.1 |
| 0.0129 | 177.0 ± 13.5 | 256.7 ± 7.8 | 36.0 ± 1.0 | 39.3 ± 4.0 | 99.7 ± 10.6 | 86.3 ± 23.5 | 350.3 ± 37.9 | 455.7 ± 14.0 | 11.0 ± 2.6 | 15.3 ± 5.5 |
| 0.041 | 189.3 ± 19.0 | 236.0 ± 25.9 | 24.0 ± 2.0 | 29.3 ± 2.1 | 94.3 ± 13.0 | 98.3 ± 6.8 | 382.3 ± 26.5 | 476.0 ± 38.7 | 8.7 ± 1.5 | 13.7 ± 2.1 |
| 0.131 | 187.7 ± 5.9 | 244.7 ± 28.0 | 29.7 ± 4.7 | 34.7 ± 6.0 | 93.7 ± 8.1 | 96.7 ± 14.2 | 303.0 ± 39.3 | 440.3 ± 52.6 | 10.7 ± 3.2 | 12.3 ± 1.5 |
| 0.415 | 185.0 ± 7.0 | 239.3 ± 13.6 | 26.3 ± 12.7 | 28.7 ± 7.1 | 101.3 ± 20.0 | 103.3 ± 9.7 | 216.3 ± 53.1 | 365.0 ± 101.4 | 11.3 ± 1.5 | 12.3 ± 2.1 |
| PC | **544.7 ± 27.2** | **612.3 ± 48.2** | **307.3 ± 11.1** | **121.7 ± 16.3** | **253.7 ± 38.5** | **880.0 ± 153.0** | **1466.0 ± 186.4** | **1730.7 ± 317.6** | **167.0 ± 52.0** | **107.3 ± 12.3** |

The results are presented as the means ± SD (N = 3). Bolded values are where IF ≥ 2 (TA97a, TA98, TA100, TA102) or IF ≥ 3 (TA1535). [a] slightly reduced bacterial background, [b] moderately reduced bacterial background. NA: not tested. Positive controls (PC): without S9: TA97a and TA98 = 4-nitroquinoline-N-oxide (0.25 µg/plate), TA100 = sodium azide (0.25 µg/plate), TA1535 = sodium azide (0.125 µg/plate), TA102 = mitomycin C (0.5 µg/plate); with S9: TA97a = benzo(a)pyrene (5 µg/plate), TA98 and TA100 = benzo(a)pyrene (2.5 µg/plate), TA102 = 2-aminoanthracene (10 µg/plate), TA1535 = 2-aminoanthracene (5 µg/plate)

### Micronucleus assay in TK6 cells

#### 24h without metabolic activation (-S9)

##### AOH

Table S 2:Comprehensive data from the CBMN assay in TK6 cells for alternariol (AOH) exposed for 24h without metabolic activation (-S9). MN: Micronuclei/Micronucleated, DMSO: dimethylsulfoxide (solvent control), MMS: methyl methane sulfonate (positive control). ns: non-significant, **: p-value <0.01, ****: p-value < 0.0001.

| AOH 24h -S9 | **1% DMSO** | **1 µM** | **2.5 µM** | **4 µM** | **5.5 µM** | **7 µM** | **8.5 µM** | **10 µM** | **MMS** |
|---|---|---|---|---|---|---|---|---|---|
| **% Survival** | 100.0 | 101.8 | 100.0 | 101.0 | 102.7 | 95.8 | 96.0 | 94.3 | 94.3 |
| **Cells with MN** | 138 | 177 | 119 | 132 | 174 | 208 | 285 | 316 | 153 |
| **Cells without MN** | 10237 | 9519 | 9995 | 9966 | 9109 | 9970 | 7950 | 7413 | 4640 |
| **% MN Cells** | 1.3 | 1.8 | 1.2 | 1.3 | 1.9 | 2.0 | 3.5 | 4.1 | 3.2 |
| **% MN** | 1.4 | 2.0 | 1.3 | 1.4 | 2.0 | 2.2 | 3.7 | 4.3 | 3.5 |
| **Fisher exact test** | **p-value** | **0.0053** | **0.3465** | **0.9026** | **0.0024** | **<0.0001** | **<0.0001** | **<0.0001** | **<0.0001** |
| | | ** | **ns** | **ns** | ** | **** | **** | **** | **** |
| **Linear Chi-square** | **p-value** | **<0.0001** | | | | | | | |
| | | **** | | | | | | | |

### AME

Table S 3: Comprehensive data from the CBMN assay in TK6 cells for alternariol monomethylether (AME) exposed for 24h without metabolic activation (-S9). MN: Micronuclei/Micronucleated, DMSO: dimethylsulfoxide (solvent control), MMS: methyl methane sulfonate (positive control). ns: non-significant, *: p-value <0.05 , ***: p-value <0.001, ****: p-value <0.0001.

| AME 24h -S9 | **1% DMSO** | **2.5 µM** | **5 µM** | **7.5 µM** | **10 µM** | **15 µM** | **17.5 µM** | **20 µM** | **MMS** |
|---|---|---|---|---|---|---|---|---|---|
| **% Survival** | 100.0 | 98.4 | 94.1 | 92.5 | 90.5 | 76.8 | 72.0 | 74.3 | 90.8 |
| **Cells with MN** | 80 | 111 | 125 | 125 | 136 | 207 | 155 | 152 | 153 |
| **Cells without MN** | 10464 | 10046 | 9992 | 10054 | 5005 | 4159 | 2663 | 2359 | 4886 |
| **% MN Cells** | 0.8 | 1.1 | 1.2 | 1.2 | 2.6 | 4.7 | 5.5 | 6.0 | 3.0 |
| **% MN** | 0.8 | 1.3 | 1.3 | 1.3 | 2.9 | 4.9 | 5.8 | 6.7 | 3.2 |
| **Fisher's exact test** | **p-value** | **0.0133** | **0.0006** | **0.0007** | **<0.0001** | **<0.0001** | **<0.0001** | **<0.0001** | **<0.0001** |
| | | * | *** | *** | **** | **** | **** | **** | **** |
| **Linear Chi-square** | **p-value** | **<0.0001** | | | | | | | |
| | | **** | | | | | | | |

<u>ALT</u>

Table S 4: Comprehensive data from the CBMN assay in TK6 cells for altenuene (ALT) exposed for 24h without metabolic activation (-S9). MN: Micronuclei/Micronucleated, DMSO: dimethylsulfoxide (solvent control), MMS: methyl methane sulfonate (positive control). ns: non-significant, *: p-value <0.05 , ****: p-value < 0.0001.

| ALT 24h-S9 | **0.5% DMSO** | **0.5 µM** | **1.25 µM** | **2.5 µM** | **3.75 µM** | **5 µM** | **12.5 µM** | **25 µM** | **37.5 µM** | **50 µM** | **100 µM** | **MMS** |
|---|---|---|---|---|---|---|---|---|---|---|---|---|
| **% Survival** | 100.0 | 99.9 | 98.0 | 92.1 | 94.8 | 94.3 | 99.2 | 91.3 | 88.1 | 95.2 | 97.0 | 94.5 |
| **Cells with MN** | 122 | 51 | 151 | 125 | 107 | 114 | 105 | 120 | 125 | 99 | 101 | 116 |
| **Cells without MN** | 10654 | 5022 | 10124 | 10179 | 10112 | 10274 | 9983 | 10310 | 10187 | 10123 | 10120 | 5022 |
| **% MN Cells** | 1.1 | 1.0 | 1.5 | 1.2 | 1.0 | 1.1 | 1.0 | 1.1 | 1.2 | 1.0 | 1.0 | 2.3 |
| **% MN** | 1.3 | 1.1 | 1.6 | 1.4 | 1.1 | 1.2 | 1.1 | 1.3 | 1.3 | 1.0 | 1.0 | 2.3 |
| **Fisher's exact test** | **p-value** | **0.5126** | **0.0328** | **0.6088** | **0.595** | **0.8444** | **0.5482** | **0.9485** | **0.6088** | **0.2509** | **0.4924** | **< 0.0001** |
| | | **ns** | * | **ns** | **ns** | **ns** | **ns** | **ns** | **ns** | **ns** | **ns** | **** |
| **Linear Chi-square** | **p-value** | **0.2448** | | | | | | | | | | |
| | | **ns** | | | | | | | | | | |

#### ATX-I

Table S 5: Comprehensive data from the CBMN assay in TK6 cells for altertoxin-I (ATX-I) exposed for 24h without metabolic activation (-S9). MN: Micronuclei/Micronucleated, DMSO: dimethylsulfoxide (solvent control), MMS: methyl methane sulfonate (positive control). ns: non-significant, *: p-value <0.05 , ***: p-value <0.001, ****: p-value < 0.0001.

| ATX-I 24h -S9 | **1% DMSO** | **0.05 µM** | **0.10 µM** | **0.21 µM** | **0.41 µM** | **1.04 µM** | **2.07 µM** | **4.15 µM** | **8.3 µM** | **MMS** |
|---|---|---|---|---|---|---|---|---|---|---|
| **% Survival** | 100.0 | 96.4 | 91.1 | 94.4 | 95.4 | 92.0 | 84.7 | 91.3 | 87.5 | 94.0 |
| **Cells with MN** | 98 | 111 | 105 | 127 | 171 | 172 | 134 | 202 | 154 | 187 |
| **Cells without MN** | 10305 | 10052 | 10161 | 10085 | 9882 | 10034 | 7016 | 9938 | 9963 | 5046 |
| **% MN Cells** | 0.9 | 1.0 | 1.0 | 1.2 | 1.7 | 1.7 | 1.9 | 2.0 | 1.5 | 3.6 |
| **% MN** | 1.0 | 1.3 | 1.1 | 1.3 | 1.8 | 1.8 | 2.1 | 2.2 | 1.6 | 3.9 |
| **Fisher's exact test** | **p-value** | **0.2973** | **0.5731** | **0.0378** | **<0.0001** | **0.0004** | **<0.0001** | **<0.0001** | **0.0002** | **<0.0001** |
| | | **ns** | **ns** | * | **** | *** | **** | **** | *** | **** |
| **Linear Chi-square** | **p-value** | **<0.0001** | | | | | | | | |
| | | **** | | | | | | | | |

### TeA

Table S 6: Comprehensive data from the CBMN assay in TK6 cells for tenuazonic acid (TeA) exposed for 24h without metabolic activation (-S9). MN: Micronuclei/Micronucleated, DMSO: dimethylsulfoxide (solvent control), MMS: methyl methane sulfonate (positive control). ns: non-significant, *: p-value <0.05 , ****: p-value < 0.0001.

| TeA 24h -S9 | **1% DMSO** | **10 µM** | **25 µM** | **50 µM** | **75 µM** | **100 µM** | **250 µM** | **500 µM** | **MMS** |
|---|---|---|---|---|---|---|---|---|---|
| **% Survival** | 100.0 | 109.4 | 100.1 | 106.5 | 102.5 | 99.0 | 98.0 | 95.2 | 101.6 |
| **Cells with MN** | 100 | 85 | 95 | 83 | 93 | 97 | 132 | 179 | 172 |
| **Cells without MN** | 10606 | 10211 | 10148 | 10175 | 10045 | 10298 | 9956 | 8170 | 5151 |
| **% MN Cells** | 0.9 | 0.8 | 0.9 | 0.8 | 0.9 | 0.9 | 1.3 | 2.1 | 3.2 |
| **% MN** | 1.0 | 0.9 | 1.0 | 0.9 | 1.0 | 1.0 | 1.4 | 2.3 | 3.6 |
| **Fisher's exact test** | **p-value** | **0.4171** | **>0.9999** | **0.3355** | **0.9424** | **>0.9999** | **0.0119** | **<0.0001** | **<0.0001** |
| | | **ns** | **ns** | **ns** | **ns** | **ns** | * | **** | **** |
| **Linear Chi-square** | **p-value** | **<0.0001** | | | | | | | |
| | | **** | | | | | | | |

#### TEN

Table S 7: Comprehensive data from the CBMN assay in TK6 cells for tentoxin (TEN) exposed for 24h without metabolic activation (-S9). MN: Micronuclei/Micronucleated, DMSO: dimethylsulfoxide (solvent control), MMS: methyl methane sulfonate (positive control). ns: non-significant, *: p-value <0.05 , ****: p-value < 0.0001.

| TEN 24h -S9 | **0.5% DMSO** | **1 µM** | **2.5 µM** | **5 µM** | **10 µM** | **25 µM** | **50 µM** | **75 µM** | **100 µM** | **200 µM** | **MMS** |
|---|---|---|---|---|---|---|---|---|---|---|---|
| **% Survival** | 100.0 | 96.7 | 100.3 | 94.5 | 102.0 | 97.9 | 96.3 | 99.1 | 101.4 | 94.9 | 71.4 |
| **Cells with MN** | 93 | 72 | 112 | 82 | 90 | 103 | 100 | 66 | 64 | 67 | 112 |
| **Cells without MN** | 10785 | 10116 | 10187 | 10344 | 10188 | 10313 | 10271 | 10268 | 10318 | 10237 | 2450 |
| **% MN Cells** | 0.9 | 0.7 | 1.1 | 0.8 | 0.9 | 1.0 | 1.0 | 0.6 | 0.6 | 0.7 | 4.4 |
| **% MN** | 0.9 | 0.8 | 1.2 | 0.8 | 0.9 | 1.1 | 1.0 | 0.7 | 0.6 | 0.7 | 4.7 |
| **Fisher's exact test** | **p-value** | **0.241** | **0.0919** | **0.5958** | **0.8822** | **0.3156** | **0.4265** | **0.0795** | **0.0451** | **0.0953** | **< 0.0001** |
| | | **ns** | **ns** | **ns** | **ns** | **ns** | **ns** | **ns** | ***** | **ns** | ******** |

### 3h with metabolic activation (+S9)

#### AOH

Table S 8: Comprehensive data from the CBMN assay in TK6 cells for alternariol (AOH) exposed for 3h with metabolic activation (+S9). MN: Micronuclei/Micronucleated, DMSO: dimethylsulfoxide (solvent control), AFB1: aflatoxin B1 (positive control). ns: non-significant, *: p-value <0.05 , ****: p-value < 0.0001.

| AOH 3h +S9 | **0.5% DMSO** | **1 µM** | **2.5 µM** | **5 µM** | **7.5 µM** | **10 µM** | **25 µM** | **50 µM** | **75 µM** | **100 µM** | **AFB1** |
|---|---|---|---|---|---|---|---|---|---|---|---|
| **%Survival** | | | | | | | | | | | |
| **Cells with MN** | 185 | 158 | 157 | 193 | 148 | 182 | 253 | 347 | 376 | 304 | 175 |
| **Cells without MN** | 10188 | 8475 | 8468 | 8408 | 7713 | 8793 | 7985 | 6105 | 4649 | 3985 | 4905 |
| **% MN Cells** | 1.78 | 1.83 | 1.82 | 2.24 | 1.88 | 2.03 | 3.07 | 5.38 | 7.48 | 7.09 | 3.44 |
| **% MN** | 2.01 | 2.03 | 1.94 | 2.55 | 2.07 | 2.22 | 3.41 | 5.97 | 8.30 | 7.72 | 3.82 |
| **Fisher's exact test** | **p-value** | **0.8269** | **0.8695** | **0.0248** | **0.6551** | **0.2243** | **<0.0001** | **<0.0001** | **<0.0001** | **<0.0001** | **<0.0001** |
| | | **ns** | **ns** | * | **ns** | **ns** | **** | **** | **** | **** | **** |
| **Linear Chi-square** | **p-value** | **<0.0001** | | | | | | | | | |
| | | **** | | | | | | | | | |

### AME

Table S 9: Comprehensive data from the CBMN assay in TK6 cells for alternariol monomethyl ether (AME) exposed for 3h with metabolic activation (+S9). MN: Micronuclei/Micronucleated, DMSO: dimethylsulfoxide (solvent control), AFB1: aflatoxin B1 (positive control). ns: non-significant, *: p-value <0.05 , **: p-value <0.01, ***: p-value <0.001, ****: p-value <0.0001

| AME 3h +S9 | **0.5% DMSO** | **2.5 µM** | **5 µM** | **7.5 µM** | **10 µM** | **25 µM** | **50 µM** | **75 µM** | **100 µM** | **125 µM** | **AFB1** |
|---|---|---|---|---|---|---|---|---|---|---|---|
| **% Survival** | 100.0 | 96.4 | 98.4 | 101.9 | 98.8 | 98.2 | 88.6 | 84.7 | 80.1 | 56.3 | 52.7 |
| **Cells with MN** | 152 | 185 | 176 | 147 | 185 | 237 | 367 | 328 | 403 | 206 | 177 |
| **Cells without MN** | 10218 | 11761 | 10427 | 10154 | 12238 | 11839 | 10943 | 13481 | 11113 | 10851 | 6593 |
| **% MN Cells** | 1.5 | 1.6 | 1.7 | 1.4 | 1.5 | 2.0 | 3.2 | 2.4 | 3.5 | 1.9 | 2.6 |
| **% MN** | 1.6 | 1.7 | 1.9 | 1.6 | 1.6 | 2.1 | 3.5 | 2.5 | 3.7 | 2.0 | 2.9 |
| **Fisher's exact test** | **p-value** | **0.6209** | **0.2659** | **0.8613** | **0.9123** | **0.0047** | **<0.0001** | **<0.0001** | **<0.0001** | **0.025** | **0.0004** |
| | | **ns** | **ns** | **ns** | **ns** | ** | **** | **** | **** | * | *** |
| **Linear Chi-square** | **p-value** | **<0.0001** | | | | | | | | | |
| | | **** | | | | | | | | | |

### ALT

Table S 10: Comprehensive data from the CBMN assay in TK6 cells for altenuene (ALT) exposed for 3h with metabolic activation (+S9). MN: Micronuclei/Micronucleated, DMSO: dimethylsulfoxide (solvent control), AFB1: aflatoxin B1 (positive control). ns: non-significant, *: p-value <0.05 , **: p-value <0.01, ****: p-value < 0.0001.

| ALT 3h +S9 | **0.5% DMSO** | **0.5 µM** | **1.25 µM** | **2.5 µM** | **3.75 µM** | **5 µM** | **12.5 µM** | **25 µM** | **37.5 µM** | **50 µM** | **AFB1** |
|---|---|---|---|---|---|---|---|---|---|---|---|
| **% Survival** | 100.0 | 97.5 | 98.5 | 99.2 | 100.5 | 96.3 | 99.2 | 103.2 | 100.8 | 99.7 | 48.4 |
| **Cells with MN** | 98 | 106 | 96 | 88 | 129 | 84 | 99 | 143 | 104 | 84 | 103 |
| **Cells without MN** | 10211 | 10114 | 10019 | 10033 | 10067 | 10148 | 10111 | 10075 | 10144 | 10115 | 4966 |
| **% MN Cells** | 1.0 | 1.0 | 0.9 | 0.9 | 1.3 | 0.8 | 1.0 | 1.4 | 1.0 | 0.8 | 2.0 |
| **% MN** | 1.0 | 1.1 | 1.0 | 0.91 | 1.4 | 0.9 | 1.0 | 1.5 | 1.1 | 0.9 | 2.1 |
| **Fisher's exact test** | **p-value** | **0.5736** | **>0.9999** | **0.5563** | **0.0327** | **0.3337** | **0.943** | **0.0029** | **0.6716** | **0.3344** | **<0.0001** |
| | | **ns** | **ns** | **ns** | * | **ns** | **ns** | ** | **ns** | **ns** | **** |
| **Linear Chi-square** | **p-value** | **0.98** | | | | | | | | | |
| | | **ns** | | | | | | | | | |

### ATX-I

Table S 11: Comprehensive data from the CBMN assay in TK6 cells for altertoxin-I (ATX-I) exposed for 3h with metabolic activation (+S9). MN: Micronuclei/Micronucleated, DMSO: dimethylsulfoxide (solvent control), AFB1: aflatoxin B1 (positive control). ns: non-significant, ****: p-value < 0.0001.

| | **0.5% DMSO** | **1 µM** | **2.5 µM** | **5 µM** | **7.5 µM** | **10 µM** | **25 µM** | **50 µM** | **75 µM** | **100 µM** | **AFB1** |
|---|---|---|---|---|---|---|---|---|---|---|---|
| **% Survival** | 100.0 | 97.3 | 99.5 | 98.1 | 98.3 | 95.8 | 86.0 | 69.0 | 55.4 | 41.4 | 54.2 |
| **Cells with MN** | 131 | 126 | 142 | 105 | 135 | 146 | 139 | 106 | 121 | 119 | 256 |
| **Cells without MN** | 10127 | 9607 | 9883 | 10011 | 10077 | 9972 | 9974 | 10027 | 10104 | 9816 | 7184 |
| **% MN Cells** | 1.3 | 1.3 | 1.4 | 1.0 | 1.3 | 1.4 | 1.4 | 1.1 | 1.2 | 1.2 | 3.4 |
| **% MN** | 1.3 | 1.4 | 1.5 | 1.1 | 1.4 | 1.6 | 1.4 | 1.1 | 1.2 | 1.3 | 3.8 |
| **Fisher's exact test** | **p-value** | **0.9499** | **0.3943** | **0.1162** | **0.8051** | **0.3331** | **0.5814** | **0.1329** | **0.5686** | **0.656** | **<0.0001** |
| | | **ns** | **ns** | **ns** | **ns** | **ns** | **ns** | **ns** | **ns** | **ns** | **** |

### TeA

Table S 12: Comprehensive data from the CBMN assay in TK6 cells for tenuazonic acid (TeA) exposed for 3h with metabolic activation (+S9). MN: Micronuclei/Micronucleated, DMSO: dimethylsulfoxide (solvent control), AFB1: aflatoxin B1 (positive control). ns: non-significant, *: p-value <0.05, ****: p-value < 0.0001.

| TeA 3h +S9 | **0,5% DMSO** | **10 µM** | **25 µM** | **50 µM** | **75 µM** | **100 µM** | **250 µM** | **500 µM** | **750 µM** | **1000 µM** | **AFB1** |
|---|---|---|---|---|---|---|---|---|---|---|---|
| **% Survival** | 100.0 | 99.4 | 98.9 | 98.2 | 94.3 | 91.6 | 85.5 | 87.0 | 85.4 | 85.8 | 49.2 |
| **Cells with MN** | 76 | 86 | 82 | 70 | 80 | 74 | 81 | 106 | 83 | 86 | 105 |
| **Cells without MN** | 10213 | 10071 | 10087 | 10081 | 10059 | 10212 | 10061 | 10059 | 10182 | 10028 | 4993 |
| **% MN Cells** | 0.7 | 0.9 | 0.8 | 0.7 | 0.8 | 0.7 | 0.8 | 1.0 | 0.8 | 0.9 | 2.1 |
| **% MN** | 0.8 | 1.0 | 0.9 | 0.7 | 0.9 | 0.8 | 0.9 | 1.1 | 0.9 | 0.9 | 2.2 |
| **Fisher's exact test** | **p-value** | **0.3869** | **0.6319** | **0.6791** | **0.6886** | **0.9347** | **0.6317** | **0.021** | **0.5783** | **0.3861** | **<0,0001** |
| | | **ns** | **ns** | **ns** | **ns** | **ns** | **ns** | * | **ns** | **ns** | **** |
| **Linear Chi-square** | **p-value** | **0.1547** | | | | | | | | | |
| | | **ns** | | | | | | | | | |

### TEN

Table S 13: Comprehensive data from the CBMN assay in TK6 cells for tentoxin (TEN) exposed for 3h with metabolic activation (+S9). MN: Micronuclei/Micronucleated, DMSO: dimethylsulfoxide (solvent control), AFB1: aflatoxin B1 (positive control). ns: non-significant, *: p-value <0.05, ****: p-value < 0.0001.

| TEN 3h +S9 | **0.5% DMSO** | **1 µM** | **2.5 µM** | **5 µM** | **10 µM** | **25 µM** | **50 µM** | **75 µM** | **100 µM** | **AFB1** |
|---|---|---|---|---|---|---|---|---|---|---|
| **% Survival** | 100.0 | 100.6 | 102.4 | 98.7 | 102.7 | 98.0 | 107.1 | 103.0 | 98.0 | 79.2 |
| **Cells with MN** | 142 | 118 | 150 | 212 | 191 | 140 | 127 | 178 | 102 | 119 |
| **Cells without MN** | 10137 | 9929 | 9174 | 13672 | 12417 | 13580 | 10796 | 10599 | 10109 | 5012 |
| **% MN Cells** | 1.4 | 1.2 | 1.6 | 1.5 | 1.5 | 1.0 | 1.2 | 1.6 | 1.0 | 2.3 |
| **% MN** | 1.5 | 1.3 | 1.7 | 1.6 | 1.6 | 1.1 | 1.3 | 1.9 | 1.0 | 2.4 |
| **Fisher's exact test** | **p-value** | **0.1906** | **0.1944** | **0.3579** | **0.4059** | **0.011** | **0.1584** | **0.1147** | **0.012** | **<0.0001** |
| | | **ns** | **ns** | **ns** | **ns** | * | **ns** | **ns** | * | **** |

### 3h without metabolic activation (-S9)

#### ALT

Table S 14: Comprehensive data from the CBMN assay in TK6 cells for altenuene (ALT) exposed for 3h without metabolic activation (-S9). MN: Micronuclei/Micronucleated, DMSO: dimethylsulfoxide (solvent control), EMS: ethyl methane sulfonate (positive control). ns: non-significant, *: p-value <0.05, ****: p-value < 0.0001.

| ALT 3h -S9 | **0.5% DMSO** | **1 µM** | **2.5 µM** | **5 µM** | **10 µM** | **25 µM** | **50 µM** | **75 µM** | **100 µM** | **200 µM** | **EMS** |
|---|---|---|---|---|---|---|---|---|---|---|---|
| **% Survival** | 100.0 | 95.1 | 92.2 | 93.3 | 95.8 | 96.3 | 91.2 | 93.6 | 94.5 | 95.9 | 38.3 |
| **Cells with MN** | 121 | 134 | 111 | 139 | 161 | 149 | 118 | 208 | 128 | 138 | 185 |
| **Cells without MN** | 9981 | 9425 | 10128 | 13858 | 10098 | 9967 | 9996 | 14021 | 10038 | 9937 | 4936 |
| **% MN Cells** | 1.2 | 1.4 | 1.1 | 1.0 | 1.6 | 1.5 | 1.2 | 1.5 | 1.3 | 1.4 | 3.6 |
| **% MN** | 1.3 | 1.6 | 1.1 | 1.1 | 1.7 | 1.6 | 1.2 | 1.6 | 1.3 | 1.5 | 3.8 |
| **Fisher's exact test** | **p-value** | **0.208** | **0.4679** | **0.1299** | **0.0263** | **0.0979** | **0.8455** | **0.0809** | **0.7025** | **0.2881** | **<0.0001** |
| | | **ns** | **ns** | **ns** | * | **ns** | **ns** | **ns** | **ns** | **ns** | **** |
| **Linear Chi-square** | **p-value** | **0.2104** | | | | | | | | | |
| | | **ns** | | | | | | | | | |

### TEN

Table S 15: Comprehensive data from the CBMN assay in TK6 cells for tentoxin (TEN) exposed for 3h without metabolic activation (-S9). MN: Micronuclei/Micronucleated, DMSO: dimethylsulfoxide (solvent control), EMS: ethyl methane sulfonate (positive control). ns: non-significant, *: p-value <0.05, **: p-value <0.01, ***: p-value <0.001, ****: p-value < 0.0001

| TEN 3h -S9 | **0.5% DMSO** | **1 µM** | **2.5 µM** | **5 µM** | **10 µM** | **25 µM** | **50 µM** | **75 µM** | **100 µM** | **200 µM** | **EMS** |
|---|---|---|---|---|---|---|---|---|---|---|---|
| **% Survival** | 100.0 | 99.9 | 96.5 | 96.2 | 100.1 | 102.0 | 98.1 | 100.5 | 96.7 | 99.6 | 36.0 |
| **Cells with MN** | 95 | 69 | 129 | 73 | 80 | 132 | 192 | 131 | 131 | 116 | 175 |
| **Cells without MN** | 10227 | 9998 | 9913 | 10084 | 10033 | 13474 | 13785 | 9984 | 9971 | 10020 | 6893 |
| **% MN Cells** | 0.92 | 0.69 | 1.28 | 0.72 | 0.79 | 0.97 | 1.37 | 1.30 | 1.30 | 1.14 | 2.48 |
| **% MN** | 0.98 | 0.74 | 1.36 | 0.78 | 0.87 | 1.02 | 1.48 | 1.34 | 1.32 | 1.22 | 2.52 |
| **Fisher's exact test** | **p-value** | **0.071** | **0.0129** | **0.1212** | **0.3245** | **0.7365** | **0.0012** | **0.011** | **0.0109** | **0.1277** | **< 0.0001** |
| | | **ns** | * | **ns** | **ns** | **ns** | ** | * | * | **ns** | **** |
| **Linear Chi-square (R)** | **p-value** | **0.0003** | | | | | | | | | |
| | | *** | | | | | | | | | |

### Fluorescence *In situ* Hybridization (FISH) data

#### AOH

Table S 16: Comprehensive data from the FISH assay for alternariol (AOH) exposed for 24h without metabolic activation (-S9). BNC: binucleated, CENT +: centromere positive, CENT-: centromere negative, MN: Micronucleated, DMSO: dimethylsulfoxide (solvent control), MMS: methyl methane sulfonate (clastogen positive control), COLC: colchicine (aneugen positive control). ns: non-significant, ****: p-value < 0.0001, N/A: not applicable.

| AOH 24h -S9 | Negative 1 (n=2) (DMSO 1%) | Negative 2 (n=2) (DMSO 1%) | Total Negative (n=4) (DMSO 1%) | AOH 5.5 µM | AOH 7 µM | AOH 8.5 µM | AOH 10 µM | MMS | COLC |
|---|---|---|---|---|---|---|---|---|---|
| # Binucleated cells (BNC) | 3785 | 6681 | 10466 | 5790 | 7688 | 5562 | 5438 | 2053 | 3964 |
| Micronucleated cells (MN) | 28 | 66 | 94 | 71 | 110 | 126 | 152 | 61 | 88 |
| % MN BNC cells | **0.74** | **0.99** | **0.90** | **1.23** | **1.43** | **2.27** | **2.80** | **2.97** | **2.22** |
| NOK | 2 | 2 | 4 | 3 | 5 | 4 | 11 | 1 | 5 |
| MN CENT+ | 4 | 12 | 16 | 15 | 19 | 23 | 19 | 9 | 54 |
| MN CENT- | 22 | 52 | 74 | 53 | 86 | 99 | 122 | 51 | 29 |
| % CENT + | **15.4** | **18.8** | **17.8** | **22.1** | **18.1** | **18.9** | **13.5** | **15.0** | **65.1** |
| % CENT - | **84.6** | **81.3** | **82.2** | **77.9** | **81.9** | **81.1** | **86.5** | **85.0** | **34.9** |
| P-value as compared with MMS | | | **0.8235 (ns)** | **0.3678 (ns)** | **0.6715 (ns)** | **0.6791 (ns)** | **0.8248 (ns)** | **N/A** | **<0.0001 (****)** |
| P-value as compared with COLC | | | **<0.0001 (****)** | **<0.0001 (****)** | **<0.0001 (****)** | **<0.0001 (****)** | **<0.0001 (****)** | **<0.0001 (****)** | **N/A** |

### AME

Table S 17: Comprehensive data from the FISH assay for alternariol monomethyl ether (AME) exposed for 24h without metabolic activation (-S9). BNC: binucleated, CENT +: centromere positive, CENT-: centromere negative, MN: Micronucleated, DMSO: dimethylsulfoxide (solvent control), MMS: methyl methane sulfonate (clastogen positive control), COLC: colchicine (aneugen positive control). ns: non-significant, ****: p-value < 0.0001, N/A: not applicable.

| AME 24h -S9 | Negative 1 (n=2) (DMSO 1%) | Negative 2 (n=2) (DMSO 1%) | Total Negative (n=4) (DMSO 1%) | AME 7.5 µM | AME 10 µM | AME 15 µM | AME 20 µM | MMS | COLC |
|---|---|---|---|---|---|---|---|---|---|
| # Binucleated cells (BNC) | 10099 | 10192 | 20291 | 9591 | 7285 | 7079 | 3032 | 7187 | 3216 |
| Micronucleated cells (MN) | 81 | 99 | 180 | 156 | 197 | 227 | 168 | 228 | 92 |
| % MN BNC | **0.80** | **0.97** | **0.89** | **1.63** | **2.70** | **3.21** | **5.54** | **3.17** | **2.86** |
| NOK | 18 | 7 | 25 | 10 | 9 | 32 | 4 | 4 | 3 |
| MN CENT+ | 19 | 21 | 40 | 30 | 32 | 32 | 19 | 34 | 57 |
| MN CENT- | 44 | 71 | 115 | 116 | 156 | 163 | 145 | 190 | 32 |
| % CENT + | **30.2** | **22.8** | **25.8** | **20.5** | **17.0** | **16.4** | **11.6** | **15.2** | **64.0** |
| % CENT - | **69.8** | **77.2** | **74.2** | **79.5** | **83.0** | **83.6** | **88.4** | **84.8** | **36.0** |
| P-value as compared with MMS | | | **0.0122 (*)** | **0.2062 (ns)** | **0.6862 (ns)** | **0.7885 (ns)** | **0.3698 (ns)** | **N/A** | **<0.0001 (****)** |
| P-value as compared with COLC | | | **<0.0001 (****)** | **<0.0001 (****)** | **<0.0001 (****)** | **<0.0001 (****)** | **<0.0001 (****)** | **<0.0001 (****)** | **N/A** |

<u>ATX-I</u>

Table S 18: Comprehensive data from the FISH assay for altertoxin-I (ATX-I) exposed for 24h without metabolic activation (-S9). BNC: binucleated, CENT +: centromere positive, CENT-: centromere negative, MN: Micronucleated, DMSO: dimethylsulfoxide (solvent control), MMS: methyl methane sulfonate (clastogen positive control), COLC: colchicine (aneugen positive control). ns: non-significant, ****: p-value < 0.0001, N/A: not applicable.

| ATX-I 24h -S9 | Negative 1 (n=2) (DMSO 1%) | Negative 2 (n=2) (DMSO 1%) | Total Negative (n=4) (DMSO 1%) | ATX-I 0.41 µM | ATX-I 1.04 µM | ATX-I 2.07 µM | ATX-I 8.3 µM | MMS | COLC |
|---|---|---|---|---|---|---|---|---|---|
| # Binucleated cells (BNC) | 10210 | 8765 | 18975 | 9342 | 10004 | 9822 | 7502 | 2374 | 2791 |
| Micronucleated cells (MN) | 77 | 86 | 163 | 130 | 139 | 128 | 125 | 102 | 52 |
| % MN BNC | **0.75** | **0.98** | **0.86** | **1.39** | **1.39** | **1.30** | **1.67** | **4.30** | **1.86** |
| NOK | 3 | 8 | 11 | 7 | 10 | 11 | 7 | 12 | 11 |
| MN CENT+ | 15 | 16 | 31 | 23 | 23 | 24 | 17 | 14 | 28 |
| MN CENT- | 59 | 62 | 121 | 100 | 106 | 93 | 101 | 76 | 13 |
| % CENT + | **20.3** | **20.5** | **20.4** | **18.7** | **17.8** | **20.5** | **14.4** | **15.6** | **68.3** |
| % CENT - | **79.7** | **79.5** | **79.6** | **81.3** | **82.2** | **79.5** | **85.6** | **84.4** | **31.7** |
| P-value as compared with MMS | | | **0.3957 (ns)** | **0.5875 (ns)** | **0.7166 (ns)** | **0.4692 (ns)** | **0.8461 (ns)** | **N/A** | **<0.0001 (****)** |
| P-value as compared with COLC | | | **<0.0001 (****)** | **<0.0001 (****)** | **<0.0001 (****)** | **<0.0001 (****)** | **<0.0001 (****)** | **<0.0001 (****)** | **N/A** |

### TeA

Table S 19: Comprehensive data from the FISH assay for tenuazonic acid (TeA) exposed for 24h without metabolic activation (-S9). BNC: binucleated, CENT +: centromere positive, CENT-: centromere negative, MN: Micronucleated, DMSO: dimethylsulfoxide (solvent control), MMS: methyl methane sulfonate (clastogen positive control), COLC: colchicine (aneugen positive control). ns: non-significant, **: p-value <0.01, ***: p-value <0.001, ****: p-value < 0.0001, N/A: not applicable.

| TeA 24h -S9 | Negative 1 (n=2) (DMSO 1%) | Negative 2 (n=2) (DMSO 1%) | Total Negative (n=4) (DMSO 1%) | TeA 75 µM | TeA 100 µM | TeA 250 µM | TeA 500 µM | MMS | COLC |
|---|---|---|---|---|---|---|---|---|---|
| # Binucleated cells (BNC) | 5318 | 5167 | 10485 | 10077 | 8977 | 8187 | 3395 | 5685 | 1270 |
| Micronucleated cells (MN) | 40 | 79 | 119 | 75 | 97 | 111 | 55 | 155 | 44 |
| % MN BNC | **0.75** | **1.53** | **1.13** | **0.74** | **1.08** | **1.36** | **1.62** | **2.73** | **3.46** |
| NOK | 6 | 14 | 20 | 16 | 26 | 18 | 10 | 13 | 5 |
| MN CENT+ | 9 | 20 | 29 | 14 | 15 | 30 | 15 | 44 | 25 |
| MN CENT- | 25 | 45 | 70 | 45 | 56 | 63 | 30 | 98 | 14 |
| % CENT + | **26.5** | **30.8** | **29.3** | **23.7** | **21.1** | **32.3** | **33.3** | **31.0** | **64.1** |
| % CENT - | **73.5** | **69.2** | **70.7** | **76.3** | **78.9** | **67.7** | **66.7** | **69.0** | **35.9** |
| P-value as compared with MMS | | | **0.8868 (ns)** | **0.4674 (ns)** | **0.2874 (ns)** | **0.7545 (ns)** | **0.6973 (ns)** | **N/A** | **0.0002 (***)** |
| P-value as compared with COLC | | | **0.0003 (***)** | **0.0001 (***)** | **<0.0001 (****)** | **0.0010 (***)** | **0.0082 (**)** | **0.0002 (***)** | **N/A** |

### In vitro Micronucleus assay in HepG2 cells

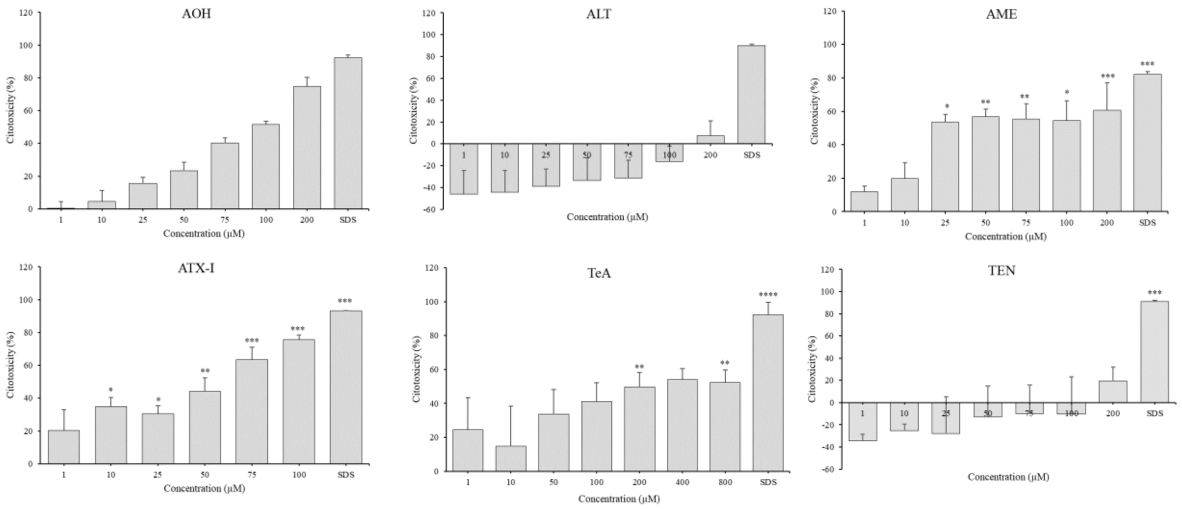


Fig S 1: Cytotoxicity of *Alternaria* toxins on HepG2 cells after 48h of incubation assessed via MTT assay. Data are normalized to untreated cells. DMSO, solvent control; SDS 0.1%, positive control. Data are presented as mean + SD (n = 3 independent experiments). *p ≤ 0.05, **p ≤ 0.01, ***p ≤ 0.001.

Table S 20: Comprehensive data from the CBMN assay in HepG2 cells for AOH.

| | BC | MNBC (‰) | NPB (‰) | NBUD (‰) | CBPI | RI (%) | Cytotoxicity (%) |
|---|---|---|---|---|---|---|---|
| Solvent control | 2000 | 11.5 ± 1.5 | 1.0 ± 0.0 | 12.5 ± 0.5 | 2.16 ± 0.10 | 100 | 0 |
| AOH (µM) | | | | | | | |
| 6.25 | 2000 | 29.5 ± 0.5 *** | 4.5 ± 0.5 | 23.0 ± 1.0 | 2.29 ± 0.03 | 114.74 ± 2.26 | -14.74 ± 2.26 |
| 12.50 | 2000 | 30.5 ± 2.5 *** | 2.0 ± 1.0 | 24.5 ± 1.5 ** | 2.08 ± 0.04 | 95.69 ± 3.96 | 4.31 ± 3.96 |
| 25.00 | 2000 | 37.5 ± 4.5 *** | 3.0 ± 0.0 | 28.0 ± 4.0 ** | 2.00 ± 0.05 | 88.90 ± 3.64 | 11.10 ± 4.02 |
| 50.00 | 2000 | 51.5 ± 2.5 *** | 5.5 ± 0.5 ** | 35.5 ± 1.5 ** | 1.64 ± 0.01 | 56.75 ± 1.26 | 43.25 ± 1.26 |
| Positive control | 2000 | 32.5 ± 0.5 *** | 3.0 ± 2.0 | 19.5 ± 0.5 | 2.25 ± 0.05 | 103.82 ± 4.77 | -10.66 ± 4.77 |

BC, Binucleated Cells; MNBC, Micronucleated Binucleated Cells; NPB, Nucleoplasmic Bridges; NBUD, Nuclear Buds; CBPI, Cytokinesis-Block Proliferation Index; RI, Replication Index; solvent control, DMSO; Positive control, Vinblastine; * $p \leq 0.05$; ** $p \leq 0.01$; *** $p \leq 0.001$

Table S 21: Comprehensive data from the CBMN assay in HepG2 cells for AME.

| | BC | MNBC (‰) | NPB (‰) | NBUD ‰ | CBPI | RI (%) | Cytotoxicity (%) |
|---|---|---|---|---|---|---|---|
| Solvent control | 2000 | 11.5 ± 1.5 | 0.5 ± 0.7 | 3.5 ± 0.7 | 1.99 ± 0.00 | 100 | 0 |
| AME (µM) | | | | | | | |
| 5.00 | 2000 | 23.5 ± 8.5 | 0.5 ± 0.7 | 6.0 ± 4.2 | 1.99 ± 0.02 | 99.90 ± 1.92 | 0.10 ± 1.92 |
| 10.00 | 2000 | 18.5 ± 1.5 | 0.0 ± 0.0 | 6.5 ± 2.1 | 1.93 ± 0.04 | 94.29 ± 4.20 | 5.71 ± 4.20 |
| 20.00 | 2000 | 27.0 ± 4.0 * | 1.5 ± 0.7 | 7.0 ± 0.0 | 1.87 ± 0.02 | 88.32 ± 2.17 | 11.68 ± 2.17 |
| 30.00 | 2000 | 29.5 ± 3.5 * | 1.5 ± 2.1 | 7.0 ± 0.5 | 1.80 ± 0.00 | 80.84 ± 0.25 | 19.16 ± 0.25 |
| Positive control | 2000 | 40.0 ± 1.5 * | 1.5 ± 0.7 | 3.5 ± 0.7 | 1.91 ± 0.01 | 91.61 ± 1.01 | 8.39 ± 1.01 |

BC, Binucleated Cells; MNBC, Micronucleated Binucleated Cells; NPB, Nucleoplasmic Bridges; NBUD, Nuclear Buds; CBPI, Cytokinesis-Block Proliferation Index; RI, Replication Index; solvent control, DMSO; Positive control, Vinblastine; * $p \leq 0.05$; ** $p \leq 0.01$; *** $p \leq 0.001$

Table S 22: Comprehensive data from the CBMN assay in HepG2 cells for ALT.

| | BC | MNBC (‰) | NPB (‰) | NBUD ‰ | CBPI | RI (%) | Cytotoxicity (%) |
|---|---|---|---|---|---|---|---|
| **Solvent control** | 2000 | 8.5 ± 0.5 | 0 | 6.0 ± 0.0 | 2.13 ± 0.04 | 100 | 0 |
| **ALT (µM)** | | | | | | | |
| **6.25** | 2000 | 13.5 ± 1.5 | 0 | 8.5 ± 2.5 | 2.07 ± 0.03 | 95.07 ± 2.57 | 4.93 ± 2.57 |
| **12.50** | 2000 | 16.5 ± 3.5 | 0 | 16.0 ± 3.0 ** | 1.96 ± 0.03 | 85.21 ± 2.57 | 14.79 ± 2.57 |
| **25.00** | 2000 | 25.5 ± 0.5 *** | 0 | 16.0 ± 1.0 ** | 1.91 ± 0.01 | 80.42 ± 0.94 | 19.58 ± 0.94 |
| **50.00** | 2000 | 27.5 ± 0.5 *** | 0 | 9.5 ± 1.5 | 1.77 ± 0.01 | 68.34 ± 0.82 | 31.66 ± 0.82 |
| **100.00** | 2000 | 34.5 ± 1.5 *** | 0 | 15.0 ± 4.0 ** | 1.68 ± 0.00 | 60.30 ± 0.13 | 39.70 ± 0.13 |
| **Positive control** | 2000 | 33.5 ± 0.5 *** | 0 | 10.5 ± 1.5 *** | 2.01 ± 0.03 | 89.25 ± 2.51 | 10.75 ± 2.51 |

BC, Binucleated Cells; MNBC, Micronucleated Binucleated Cells; NPB, Nucleoplasmic Bridges; NBUD, Nuclear Buds; CBPI, Cytokinesis-Block Proliferation Index; RI, Replication Index; solvent control, DMSO; Positive control, Vimblastine; * $p \leq 0.05$; ** $p \leq 0.01$; *** $p \leq 0.001$

Table S 23: Comprehensive data from the CBMN assay in HepG2 cells for ATX-I.

| | BC | MNBC (‰) | NPB (‰) | NBUD (‰) | CBPI | RI (%) | Cytotoxicity (%) |
|---|---|---|---|---|---|---|---|
| **Solvent control** | 2000 | 8.0 ± 0.0 | 3.5 ± 0.5 | 7.5 ± 1.5 | 2.28 ± 0.06 | 100 | 0 |
| **ATX-I (µM)** | | | | | | | |
| **0.25** | 2000 | 23.0 ± 0.0 *** | 2.0 ± 1.0 | 10.5 ± 1.5 | 2.23 ± 0.00 | 109.50 ± 0.25 | -9.50 ± 0.25 |
| **0.50** | 2000 | 28.5 ± 2.5 *** | 2.5 ± 0.5 | 12.5 ± 1.5 | 2.11 ± 0.01 | 98.98 ± 1.32 | 1.02 ± 1.32 |
| **1.00** | 2000 | 32.5 ± 0.5 *** | 1.0 ± 0.0 | 13.5 ± 1.5 | 2.07 ± 0.01 | 94.80 ± 0.57 | 5.20 ± 0.57 |
| **Positive control** | 2000 | 36.0 ± 2.0 *** | 6.0 ± 2.0 | 10.0 ± 1.0 | 2.10 ± 0.00 | 97.65 ± 0.44 | 2.35 ± 0.44 |

BC, Binucleated Cells; MNBC, Micronucleated Binucleated Cells; NPB, Nucleoplasmic Bridges; NBUD, Nuclear Buds; CBPI, Cytokinesis-Block Proliferation Index; RI, Replication Index; solvent control, DMSO; Positive control, Vinblastine; * $p \leq 0.05$; ** $p \leq 0.01$; *** $p \leq 0.001$

Table S 24: Comprehensive data from the CBMN assay in HepG2 cells for TeA.

| | BC | MNBC (‰) | NPB (‰) | NBUD ‰ | CBPI | RI | Cytotoxicity (%) |
|---|---|---|---|---|---|---|---|
| **Solvent control** | 2000 | 16.0 ± 2.0 | 1 ± 0.0 | 1 ± 0.0 | 2.06 ± 0.01 | 100 | 0 |
| **TeA (µM)** | | | | | | | |
| **1.00** | 2000 | 17.0 ± 4.0 | 0 | 2.5 ± 0.5 | 2.07 ± 0.04 | 100.94 ± 3.44 | -0.94 ± 3.44 |
| **10.00** | 2000 | 13.0 ± 0.0 | 0 | 4.5 ± 0.5 | 2.02 ± 0.02 | 96.37 ± 1.98 | 3.63 ± 1.98 |
| **50.00** | 2000 | 30.5 ± 5.5 * | 0.5 ± 0.5 | 6.5 ± 2.5 | 1.70 ± 0.03 | 65.99 ± 3.25 | 34.01 ± 3.25 |
| **75.00** | 2000 | 28.5 ± 5.5 * | 0 | 2.5 ± 0.5 | 1.44 ± 0.01 | 41.87 ± 0.52 | 58.13 ± 0.52 |
| **Positive control** | 2000 | 40.0 ± 3.0 * | 0 | 2.0 ± 1 | 2.06 ± 0.01 | 100.00 ± 0.71 | 0.00 ± 0.71 |

BC, Binucleated Cells; MNBC, Micronucleated Binucleated Cells; NPB, Nucleoplasmic Bridges; NBUD, Nuclear Buds; CBPI, Cytokinesis-Block Proliferation Index; RI, Replication Index; solvent control, DMSO; Positive control, Vinblastine; * $p \leq 0.05$; ** $p \leq 0.01$; *** $p \leq 0.001$

Table S 25: Comprehensive data from the CBMN assay in HepG2 cells for TEN.

| | BC | MNBC (‰) | NPB (‰) | NBUD (‰) | CBPI | RI (%) | Cytotoxicity (%) |
|---|---|---|---|---|---|---|---|
| **Solvent control** | 2000 | 9.0 ± 1.0 | 3.0 ± 0.0 | 4.5 ± 1.5 | 2.24 ± 0.01 | 100 | 0 |
| **TEN (µM)** | | | | | | | |
| **6.25** | 2000 | 13.5 ± 0.5 | 1.0 ± 0.0 | 8.5 ± 0.5 | 2.12 ± 0.02 | 99.02 ± 2.01 | 0.98 ± 2.01 |
| **12.50** | 2000 | 14.0 ± 0.0 | 5.5 ± 1.5 | 6.5 ± 0.5 | 2.02 ± 0.01 | 90.81 ± 0.94 | 9.19 ± 0.94 |
| **25.00** | 2000 | 18.5 ± 0.5 * | 2.5 ± 0.5 | 8.0 ± 0.0 | 2.04 ± 0.02 | 92.41 ± 2.07 | 7.59 ± 2.07 |
| **50.00** | 2000 | 19.5 ± 0.5 ** | 4.5 ± 0.5 | 13.0 ± 2.0 ** | 2.01 ± 0.04 | 89.48 ± 3.71 | 10.52 ± 3.71 |
| **100.00** | 2000 | 25.0 ± 2.0 *** | 1.0 ± 1.0 | 12.5 ± 0.5 ** | 1.87 ± 0.01 | 76.82 ± 0.63 | 23.18 ± 0.63 |
| **Positive control** | 2000 | 35.0 ± 2.0 *** | 7.5 ± 2.5 | 16.5 ± 2.5 ** | 2.05 ± 0.04 | 93.38 ± 3.45 | 6.62 ± 3.45 |

BC, Binucleated Cells; MNBC, Micronucleated Binucleated Cells; NPB, Nucleoplasmic Bridges; NBUD, Nuclear Buds; CBPI, Cytokinesis-Block Proliferation Index; RI, Replication Index; solvent control, DMSO; Positive control, Vinblastine; * $p \leq 0.05$; ** $p \leq 0.01$; *** $p \leq 0.001$

### γH2AX assay in HepaRG cells

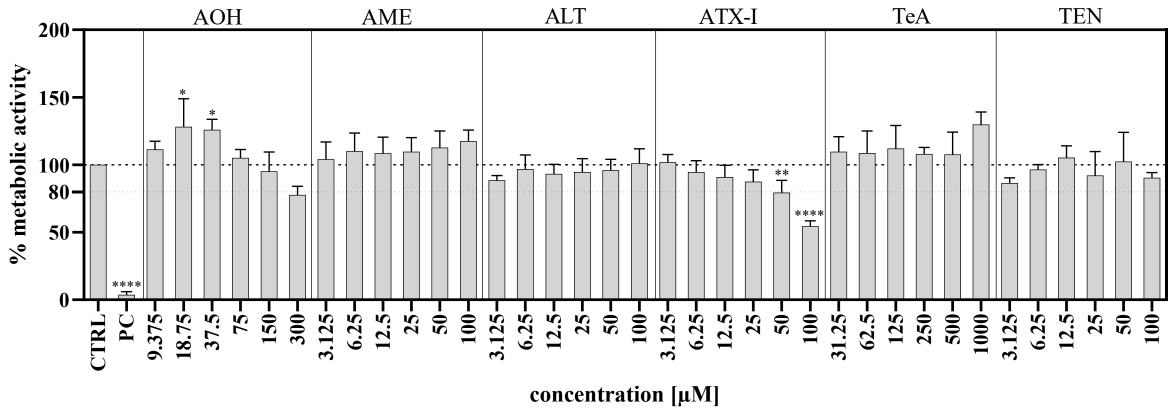


Fig S 2: Cytotoxicity of Alternaria toxins on HepaRG cells after 24h-exposure assessed by MTT assay. Data were normalized to untreated cells (CTRL). Cells exposed to 0.01% Triton-X served as positive control (PC). Data are presented as mean + SD with $^*p \leq 0.05$, $^{**}p \leq 0.01$, $^{****}p \leq 0.0001$.
